# Whole-genome sequencing of multiple isolates of *Puccinia triticina* reveals asexual lineages evolving by recurrent mutations

**DOI:** 10.1101/2020.07.22.212464

**Authors:** John P. Fellers, Sharadha Sakthikumar, Fei He, Katie McRell, Guus Bakkeren, Christina A. Cuomo, James A. Kolmer

## Abstract

**Background:** The wheat leaf rust fungus, *Puccinia triticina* Erikss. is a worldwide pathogen of tetraploid durum and hexaploid wheat. Many races of *P. triticina* differ for virulence to specific leaf rust resistance genes and are found in most wheat-growing regions of the world. Wheat cultivars with effective leaf rust resistance exert selection pressure on *P. triticina* populations for virulent race types. The objectives of this study were to examine whole-genome sequence data of 121 *P. triticina* isolates and to gain insight into race evolution. The collection included isolates comprising many different race phenotypes collected worldwide from common wheat in the U.S. and the European Union together with isolates from durum wheat. One isolate from the wild wheat relative *Aegilops speltoides,* and two from *Ae. cylindrica* were also included for comparison.

**Results:** Based on 121,907 variants identified relative to the reference race 1-1 genome, the isolates were clustered into 11 major lineages with 100% bootstrap support. The isolates were also grouped based on variation in approximately 1400 secreted resistance interactor candidate proteins. In gene-coding regions, all groups had high ratios of non-synonymous to synonymous mutations and nonsense to readthrough mutations.

**Conclusions:** Based on total variation or variation in the secreted protein genes, isolates grouped the same indicating that variants were distributed across the entire genome. Our results suggest that recurrent mutation and selection play a major role in differentiation within the clonal lineages.

## BACKGROUND

Rust diseases of cereal crops are widespread throughout grain-growing regions of the world and are capable of causing severe yield loss (Saari and Prescott 1985). Of the three rust diseases on wheat (Kolmer et al 2019), leaf rust caused by *Puccinia triticina* Erikss., is the most widespread and is the leading cause of yield loss on a worldwide basis (Savary et al 2019). In the United States, leaf rust is found in the soft red winter wheat region of the eastern and southern states, the hard red winter wheat region of the southern Great Plains and in the hard red spring wheat region of the northern Great Plains (Kolmer 2019). In 2007, a severe leaf rust epidemic occurred in the Great Plains region and losses in Kansas and Minnesota were estimated at 14% and 7%, respectively (Kolmer et al 2009).

Like the other cereal rusts, *P. triticina* is a heteroecious rust that requires two taxonomically unrelated hosts to complete the sexual cycle, and is macrocyclic with five distinct spore stages (Kolmer et al. 2019). The principle hosts of *P. triticina* are common hexaploid wheat *Triticum aestivum* L.; tetraploid durum wheat *T. turgidum* L. ssp. *durum*; and wild emmer wheat *T. turgidum ssp. dicoccoides* (Bolton et al. 2008). Infections of *P. triticina* are also found on *Aegilops cylindrica* L. (goatgrass) in the southern Great Plains region. Dikaryotic urediniospores are produced asexually on susceptible host plants and can cycle indefinitely. Dissemination of the disease occurs by wind-blown urediniospores that are deposited in wheat fields by precipitation events. The urediniospores germinate and penetrate the host plants through the stomata and form haustoria in the mesophyll cells below the epidermal and palisade layers of cells. New uredinia erupt on the upper leaf surface within 7-10 days after the initial infection. Two-celled, dikaryotic teliospores are formed in the old uredinia (subsequently called telia), which are the overwintering stage. In the spring, teliospores undergo a brief diploid phase, and then each cell germinates to produce four haploid basidiospores that infect alternate hosts. The sexual cycle is completed by the production of haploid spermagonia and spermatia on the upper leaf surface. Fertilization results in dikaryotic aeciospores in aecial cups on the lower leaf surface. The aeciospores are wind-dispersed and infect the uredinial-telial hosts. The most compatible alternate host of *P. triticina* is *Thalictrum speciosissimum* L, common meadow rue. Susceptible alternate hosts in *Thalictrum* spp. are found in southern Europe and West Asia (Jackson and Mains, 1921; Saari et al. 1968), however in North America, native *Thalictrum* species have limited susceptibility to basidiospore infection. Populations of *P. triticina* in North America (Ordonez and Kolmer 2009) and worldwide (Kolmer et al 2019) have high levels of linkage disequilibrium and higher than expected heterozygosity for microsatellite or simple sequence repeat (SSR) loci, which strongly indicated clonal reproduction.

*Puccinia triticina* interacts with wheat in a gene-for-gene manner (Samborski and Dyck 1968: 1976) as described by Flor (1942, 1971). Specific leaf rust resistance genes in wheat interact with specific effector gene products in avirulent isolates to produce incompatible infection types, which range from minute hypersensitive necrotic flecks to small uredinia that are surrounded by necrosis or chlorosis (Bolton et al. 2008). Isolates that lack corresponding effectors and are virulent to the specific leaf rust resistance gene, produce large uredinia without chlorosis or necrosis and can produce hundreds of urediniospores daily. Since the mid-1940s (Kolmer 2019) wheat breeding programs in the Great Plains region have released wheat cultivars with leaf rust resistance genes that provide resistance to most of the *P. triticina* population. However, within a few years after release, virulent genotypes of *P. triticina* arise and increase, rendering the cultivars susceptible. As a result of the constant selection for virulent genotypes, many different races of *P. triticina* are found in the U.S. (Kolmer 2019) and worldwide (Kolmer et al. 2019). Races of *P. triticina* are determined by the avirulent or virulent classification of infection types produced by single-uredinial isolates on wheat genotypes that differ by single leaf rust resistance genes (Bolton et al 2008).

In the absence of the sexual cycle, the source of new races in *P. triticina* in North America has been assumed to be somatic mutation of urediniospores. Some isolates of *P. triticina* have been shown to be heterozygous for avirulence/virulence alleles to specific leaf rust resistance genes (Samborski and Dyck, 1968, 1976; Statler 1977; 1979; 1982; 2000). Mutations at heterozygous loci can lead to loss of recognition by the host resistance protein and result in a compatible infection type (Kolmer and Dyck 1994). Infections of *P. triticina* can survive the winter in a large area of the winter wheat region of the southern U.S., resulting in a large population in which mutation and selection of virulent genotypes due to host resistance genes can occur.

A reference genome sequence of *P. triticina* was recently produced (Coumo et al. 2017). The genome was of race 1-1, designated as BBBDB in the *P. triticina* nomenclature system (Long and Kolmer, 1989). This isolate is avirulent to most leaf rust resistance genes and is heterozygous for avirulence/virulence at a number of effector loci (Samborski and Dyck, 1968). The genome contains 135 Mb of DNA, 51% mobile and repetitive elements, with an estimated 14,800 genes. The reference genome is very useful for alignment of sequence reads for identifying and assessing genetic polymorphisms in populations of *P. triticina* using whole-genome sequencing of sample isolates.

In this study, a large number of *P. triticina* isolates collected from common wheat in North America and Europe, with many different race phenotypes, were sequenced and compared with the race 1-1 BBBDB reference genome. In addition, isolates collected from durum wheat worldwide, and the wheat relatives *Ae. cylindrica* and *Ae. speltoides* were also sequenced and compared with the reference genome. The objectives of the research were: 1) to determine the phylogenetic lineage relationships in a diverse collection of *P. triticina*, 2) to characterize the genetic variation within lineages, and 3) to examine the relationship between molecular phylogeny and race phenotype.

## RESULTS

The genomes of 121 isolates of *P. triticina* were included in this study (Supplemental Table 1). Eighty-five isolates were from North America, with 65 different virulence phenotypes; 21 isolates were collected from durum wheat from various countries world-wide with three different virulence phenotypes; and 12 isolates were collected from common wheat in Europe with 11 different virulence phenotypes. A single isolate from *Ae. speltoides* was collected in Israel, and two isolates from North America were collected from *Ae. cylindrica* (common goatgrass). The virulence phenotype, country of origin, sequence statistics, avirulence/virulence formula, and references to previous characterization by earlier marker systems for each isolate is listed in Supplemental Table 1. The isolate from *Ae. speltoides* (Pt_ISR_850) had the lowest alignment to the reference genome at 78%, all other isolates had > 90% alignment. Average depth of coverage across the reference genome ranged from 9X (isolate 11US045_2 TNBGJ) to 95X (isolate 11US220_3 MFNSB).

Maximum parsimony analysis was used to determine relationships between isolates based on SNP genotypes. Using 121,907 informative SNPs from sites with <= 5% missing calls, the isolates could be clustered into eleven major groups with 100% support. The isolates from North America (NA) clustered into six discrete groups (Fig 1). The genome reference BBBDB race 1-1, collected in 1954 (Ordonez and Kolmer, 2009), was in the NA1 group with three other isolates. Isolate 04NE356, with the similar virulence phenotype BBBDJ, was collected 50 years later and differed by only 4,447 SNPs when compared to the reference genome. The NA2 group consisted of eleven isolates with ten different virulence phenotypes that were all avirulent to the isogenic *Lr2a* line, and virulent to lines with genes *Lr2c* and *Lr28* (Supplemental Table 2. The 35 NA3 group isolates consisted of 25 phenotypes that were all virulent to genes *LrB, Lr3bg, Lr17*, and avirulent to *Lr28*. These virulence phenotypes were first found in large numbers in the mid 1990’s after the release of *Lr37* in hard red winter cultivars in the southern Great Plains (Ordonez and Kolmer, 2009; Kolmer 2019). The two isolates in the NA4 group were both virulent to wheat lines with *Lr1, Lr2a, Lr2c, Lr17, Lr28*, and avirulent to *Lr3*. These isolates were originally collected from *Ae. cylindric*a in the southern Great Plains and were widely separated from the other NA groups. The NA5 group consisted of 32 isolates with 23 different virulence phenotypes that were avirulent to lines with genes *LrB*, *Lr17*, and virulent *Lr28*. Two isolates PBJL_PRTUS6 and PBJSF_84VAPA, that are both avirulent to wheat lines with *Lr2a*, and virulent to *Lr2c, Lr11* and *Lr17,* were highly related and were placed in the NA6 group. A single isolate, TCTDL_03VA190, did not cluster with isolates in any other group.

**Figure 1.**
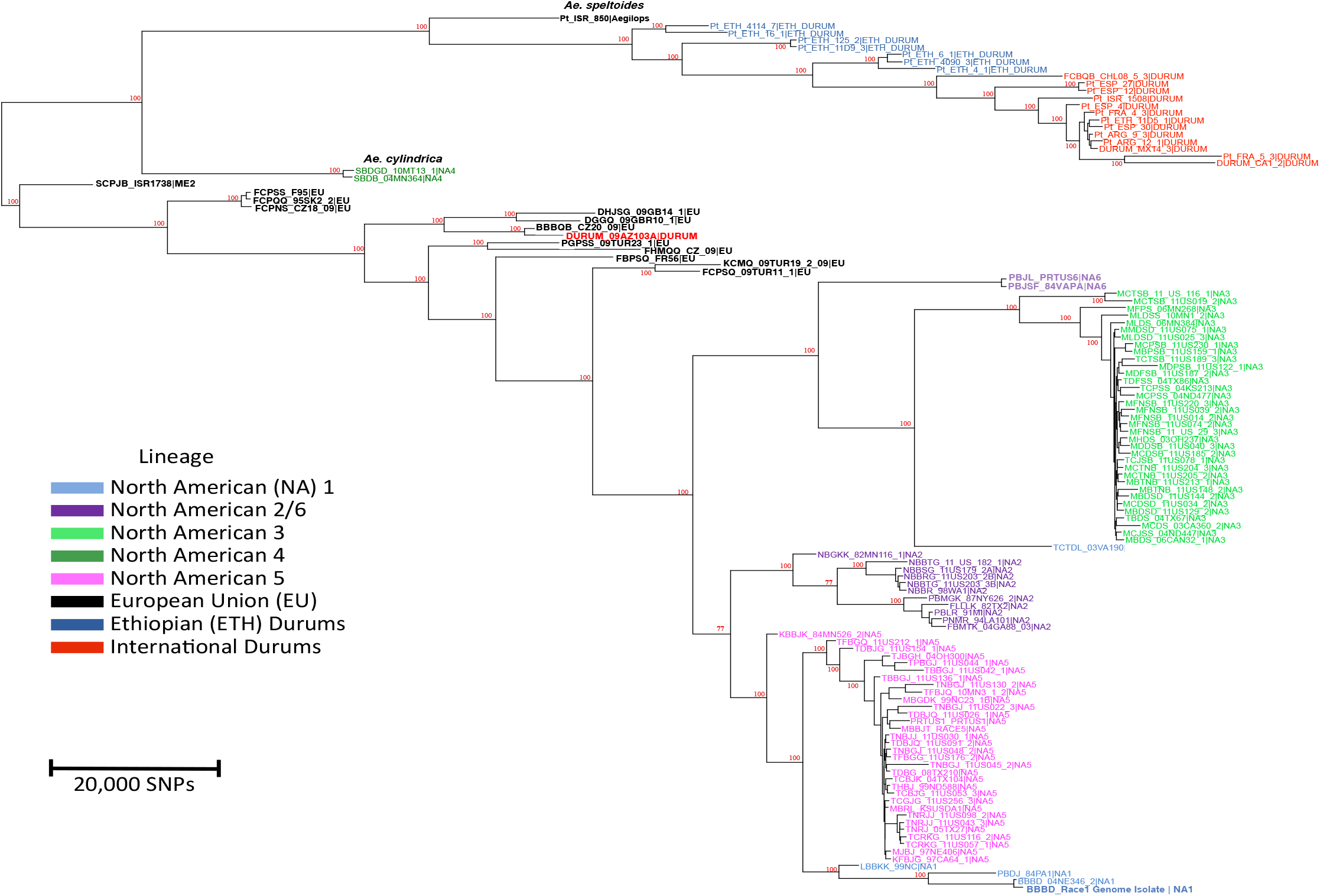
Relationship tree of *Puccinia triticina* isolates. Relatedness based on PAUP parsimony analysis and bootstrap analysis of SNPs across the genome. Total number of SNPs included was 134,940, of which 121,907 were parsimony-informative. Markers used were 95% informative at each site. Colors represent population origin of each isolate.

Thirteen isolates from Europe (EU) with 12 different virulence phenotypes were all virulent to *LrB*, avirulent to *Lr28*, and varied for virulence to the other *Lr* genes (Supplemental Table 2, Fig. 1). The European isolates were also more diverse for SNP genotypes, but were grouped discretely from all isolates in the NA groups. One isolate, DBBQJ_09AZ, was collected from durum wheat in Arizona, but was grouped with the European isolates since it was highly related to those isolates for virulence and SNP genotype. All other isolates collected from durum wheat had distinct SNP genotypes compared to the isolates from hexaploid wheat. The seven Ethiopian durum (ETH_DURUM) isolates were distinct for SNP genotype compared to the other isolates from durum. These isolates are all avirulent to cultivar Thatcher, the recurrent parent of the differential lines, and thus were not phenotyped. Isolates collected from durum wheat in California, Mexico, Argentina, Spain, Ethiopia, France, and Israel (DURUM) were closely related for SNP genotype, and were distinct from all other isolates. These isolates were all virulent to *Lr14b* and *Lr20* (Supplemental Table 2), and avirulent to most of the other Lr genes in the differential set. A single isolate collected in Israel from diploid wheat progenitor *Ae. speltoides*, had a distinct SNP genotype and did not cluster with isolates from any other group. The isolate SCPBJ_ISR173, collected in Israel from common wheat, also did not group with any other isolates in any of the genotype groups but was closest for genotype to the two isolates in NA4 that had similar virulence phenotypes.

Excluding the isolate from *Ae. speltoides*, isolates in NA4 had the lowest average number of SNPs relative to the race 1-1 reference genome at 310,033 and the durum isolates in the ETH-Durum group had the highest average number of SNPs at 450,544 (Table 1). Isolates in NA4 also had the lowest Π value of 2.29 × 10^−3^ and isolates in the ETH-durum group had the highest Π value of 3.33 × 10^−3^. The isolates from the groups collected from common wheat had higher levels of SNP heterozygosity ranging from 76% for NA3 to 97% for NA1. The International durum and Ethiopian durum groups had lower levels of SNP heterozygosity at 56% (Table 1). Even though the isolate from *Ae. speltoides* had the highest number of SNPs relative to the Race 1-1 reference genome at 996,187 only 35% were heterozygous. The large majority of the SNPs in all isolates occurred in the non-expressing intergenic regions, between 78.5% for the isolate from *Ae. speltoides* to 84.6% for isolates in NA4. The ratios of nonsynonymous to synonymous SNPs within the expressed regions were greater than 1.0 for all groups, and ratios for nonsense to read through SNPs were greater than 4.0 for all groups.

**Table 1.** Genome wide SNP diversity of *Puccinia triticina* isolates in groups from North America (NA) and Europe (EU), two groups virulent to durum wheat, and the wheat progenitor *Aegilops speltoides*

| Population | Average total SNPs | Average diversity $\pi$ (nucleotide) | Average Intergenic SNPs % | Average SNP Heterozygosity % | Average Non-synonymous/synonymous SNPs | Average Nonsense/Read-through SNPs |
| --- | --- | --- | --- | --- | --- | --- |
| NA1 | 321,326 | $2.37 \times 10^{-3}$ | 84.0 | 97 | 1.659 (21,404/12,899) | 4.237 (603/142) |
| NA2 | 422,333 | $3.12 \times 10^{-3}$ | 83.7 | 87 | 1.632 (28,383/17,389) | 4.586 (796/173) |
| NA3 | 342,861 | $2.53 \times 10^{-3}$ | 84.2 | 76 | 1.657 (22,594/13,630) | 4.311 (657/152) |
| NA4 | 310,033 | $2.29 \times 10^{-3}$ | 84.6 | 66 | 1.702 (20,426/12,000) | 4.992 (646/129) |
| NA5 | 322,011 | $2.38 \times 10^{-3}$ | 84.1 | 90 | 1.654 (21,478/12,980) | 4.345 (628/145) |
| NA6 | 346,721 | $2.56 \times 10^{-3}$ | 84.2 | 81 | 1.669 (22,974/13,765) | 4.369 (680/155) |
| Intl-Durum | 416,611 | $3.08 \times 10^{-3}$ | 84.0 | 56 | 1.645 (27,469/16,696) | 4.258 (738/174) |
| ETH-Durum | 450,544 | $3.33 \times 10^{-3}$ | 84.1 | 56 | 1.657 (29,843/18,010) | 4.254 (791/186) |
| Europe | 413,951 | $3.06 \times 10^{-3}$ | 83.8 | 81 | 1.635 (27,723/16,949) | 4.355 (794/182) |
| *Aegilops speltoides* | 996,187 | $7.36 \times 10^{-3}$ | 78.4 | 35 | 1.155 (69,440/60,101) | 4.045(1505/372) |

The isolates were also grouped in a two-dimensional PCA plot simply based on virulence to the 22 Thatcher *Lr* isolines (Figure 2). The isolates did not fall into discrete nonoverlapping clusters. However, there were regions of the PCA plot where most of the isolates were from one group. Of the NA groups, isolates in NA1, NA2, NA4, and NA5 were grouped more closely in the right side of the plot with positive first axis values, while isolates in NA3 and NA6 all had negative first axis values. Isolates in the EU group were diverse for virulence and were found in both sides of the plot. Of the NA groups with more than 10 isolates, isolates in NA3 had the smallest average virulence difference at 3.50, and isolates in NA5 had the largest at 4.85 (Table 2). Isolates in the International Durum group were very closely related for virulence, differing on average by only 1.03. The average virulence difference overall the 121 isolates was 7.90.

**Figure 2.**
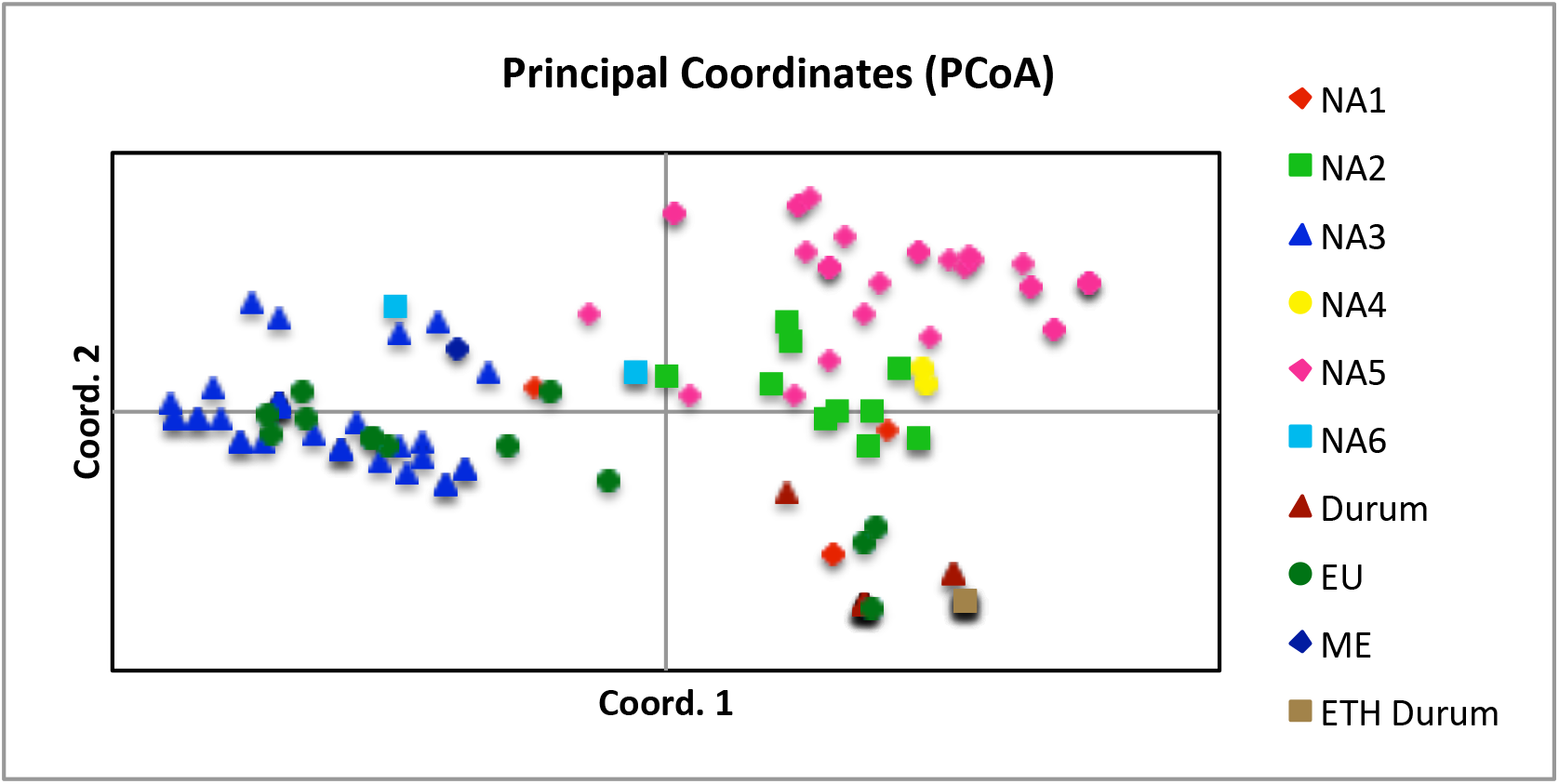
Principal coordinate plot based on virulence to 20 lines of Thatcher wheat near isogenic for leaf rust resistance genes of *Puccinia triticina* isolates in groups from North America (NA) and Europe (EU) and two groups virulent to durum wheat.

**Table 2.** Virulence diversity of Puccinia triticina isolates in SNP genotype groups from North America (NA), Europe (EU), two groups virulent to durum wheat, and the wheat progenitor *Aegilops speltoides*.

| Group | No. of isolates | No. of virulence phenotypes | Average virulence difference |
| --- | --- | --- | --- |
| NA1 | 4 | 3 | 3.30 |
| NA2 | 11 | 10 | 3.84 |
| NA3 | 35 | 25 | 3.50 |
| NA4 | 2 | 2 | 1.00 |
| NA5 | 32 | 23 | 4.85 |
| NA6 | 2 | 2.0 |  |
| EU | 13 | 12 | 4.63 |
| Intl Durum | 13 | 3 | 1.03 |
| ETH Durum | 8 | --- | --- |

The total SNP variation (intronic and intergenic) was then compared to SNPs that occurred only in predicted secreted proteins to determine if there were differences in the genetic relationship between isolates based on total genomic variation and variation that occurred in genes involved with pathogenicity and virulence specificity. Isolates in ETH-Durum and the *Ae. speltoides* isolate were not included due to their high divergence from the other isolate groups. The two-dimensional principal coordinate plot of the total 343,000 SNPs across the genome (Figure 3A) grouped the isolates in the same manner as in the neighbor joining plot. Analysis of variation in predicted secreted peptides was based on 16,000 SNPs within 2,000 bp 5’ of the start codon open reading frame, and 2,000 bp 3’ of the stop codon of ca. 1,400 genes. Isolates were clustered the same as for the total genomic data (Figure 3B). Median diversity (*π*) for all SNP’s and SNPs within secreted peptides averaged >40% (Figure 4a) indicating very large genetic variation across each contig or gene, with the total genomic SNPs being slightly more diverse, although within the standard deviation of the average diversity of the secreted peptides. The median *Fst* measures population differentiation and divergence for each contig or gene. A generally higher median (Figure 4b) and 2X higher median density (Figure 4c) was seen for secreted peptides, though within standard deviation of the total genomic mean.

**Figure 3.**
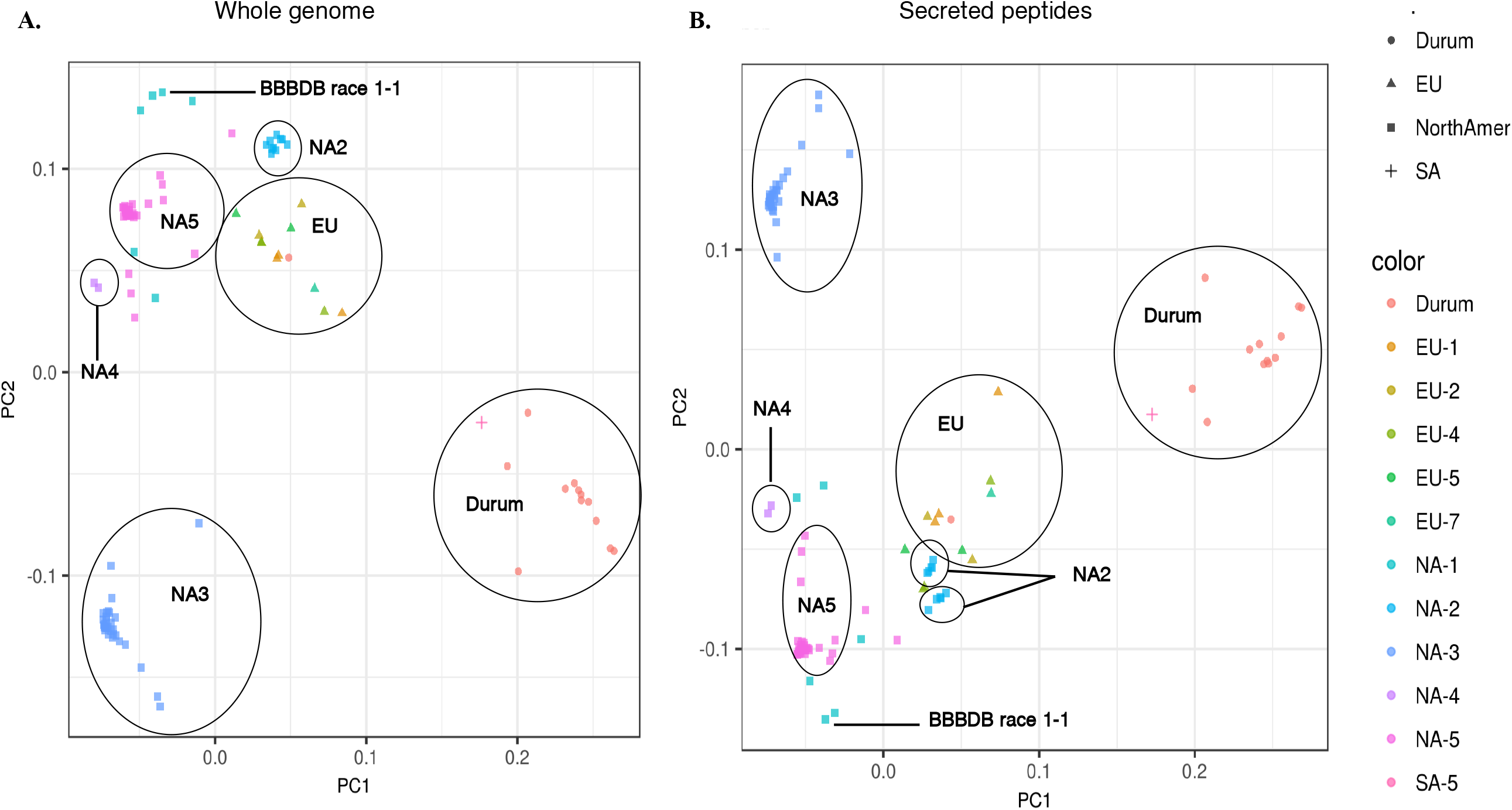
Principal component (PC) analysis of 121 *P. triticina* isolates representing populations from North America, Europe, South America and from different wheat classes and species. Data is based on A) ~343,000 whole genome SNPs, or B) ~16,000 SNPs found in secreted peptides.

**Figure 4.**
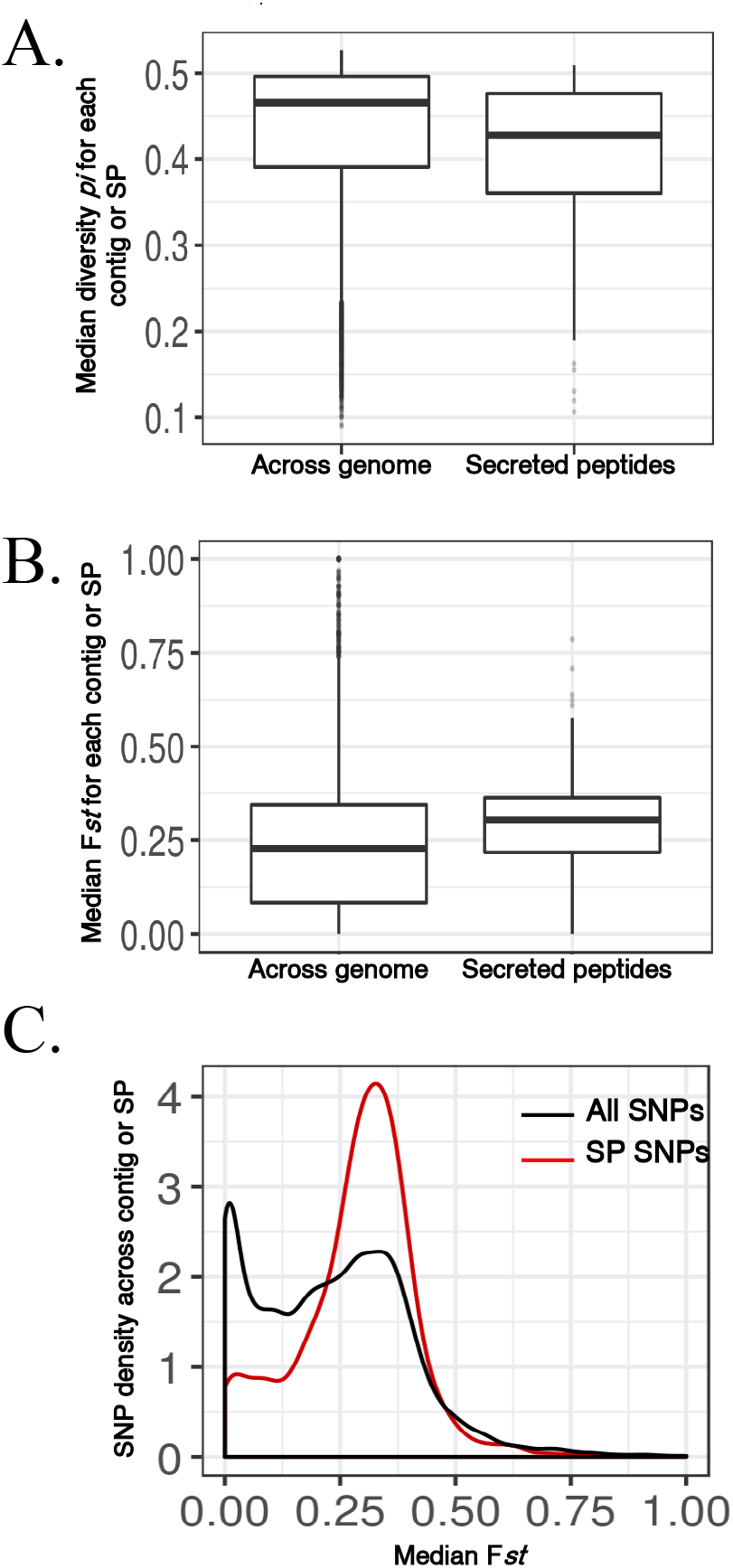
A) Median diversity (P*i*) for each contig, first for all of the SNPs, then SNPs within secreted peptides (SP). B) Median F*st* measuring population substructure and divergence for each contig, first for all of the SNPs, then SNPs within secreted peptides; C) Median F*st* for a contig versus density of SNPs across the contig or SP.

The relationship between virulence phenotype and SNP genotype was further examined. In clonal populations such as *P. triticina*, it would be expected to find at least a general correlation between phenotype and genotype. As genetic similarity between isolates increased to over 90%, the similarity of isolates for virulence phenotype increased to over 60% (Fig. 5). At this high level of genetic similarity there was no difference in phenotypic similarity between total genomic SNPs and SNPs in predicted secreted proteins.

**Figure 5.**
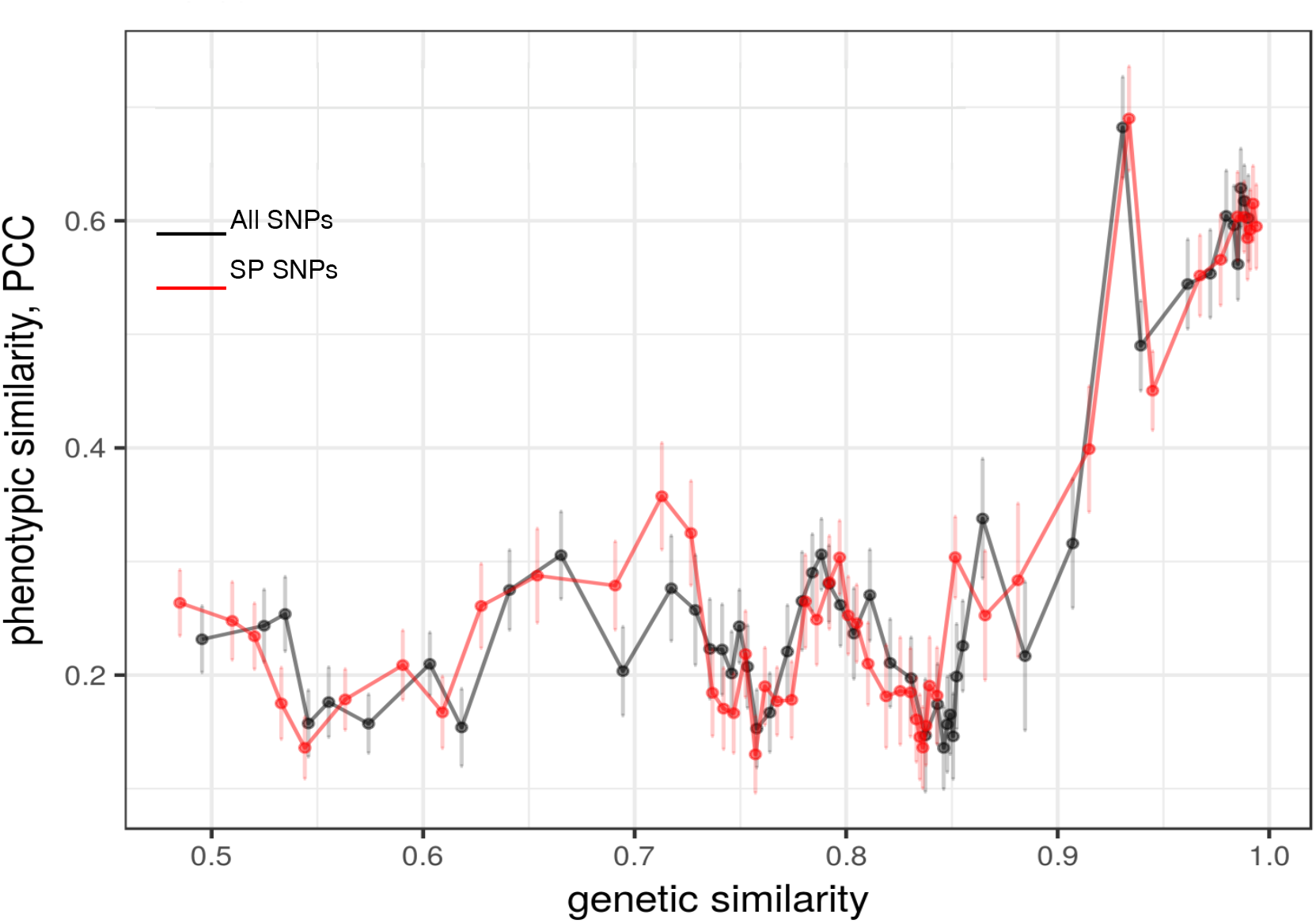
Genetic similarity as measured by identity-by-state of the 121 rust isolates based on all SNPs across the genome (black) or just SNPs within secreted peptides (red). Each data point represents five percentile bins with 95% confidence interval.

A genome-wide association study (GWAS) was conducted based on avirulence and virulence to the near isogenic Thatcher lines that differ for single resistance genes. The association analysis was conducted for total genomic SNPs and also for SNPs within secreted peptides. More than 400 contigs in the total genomic data (Figure 6a) and SNPs in 50 contigs predicted secreted peptides (Figure 6b) were associated with virulence/avirulence to Thatcher lines with *Lr17* and *Lr28*. Relatively few contigs were associated with virulence/avirulence to lines with *LrB*, *Lr10*, *Lr16*, and *Lr21*. Virulence/avirulence to nine of the Thatcher lines was associated with less than 15 SNPs in predicted secreted proteins.

**Figure 6.**
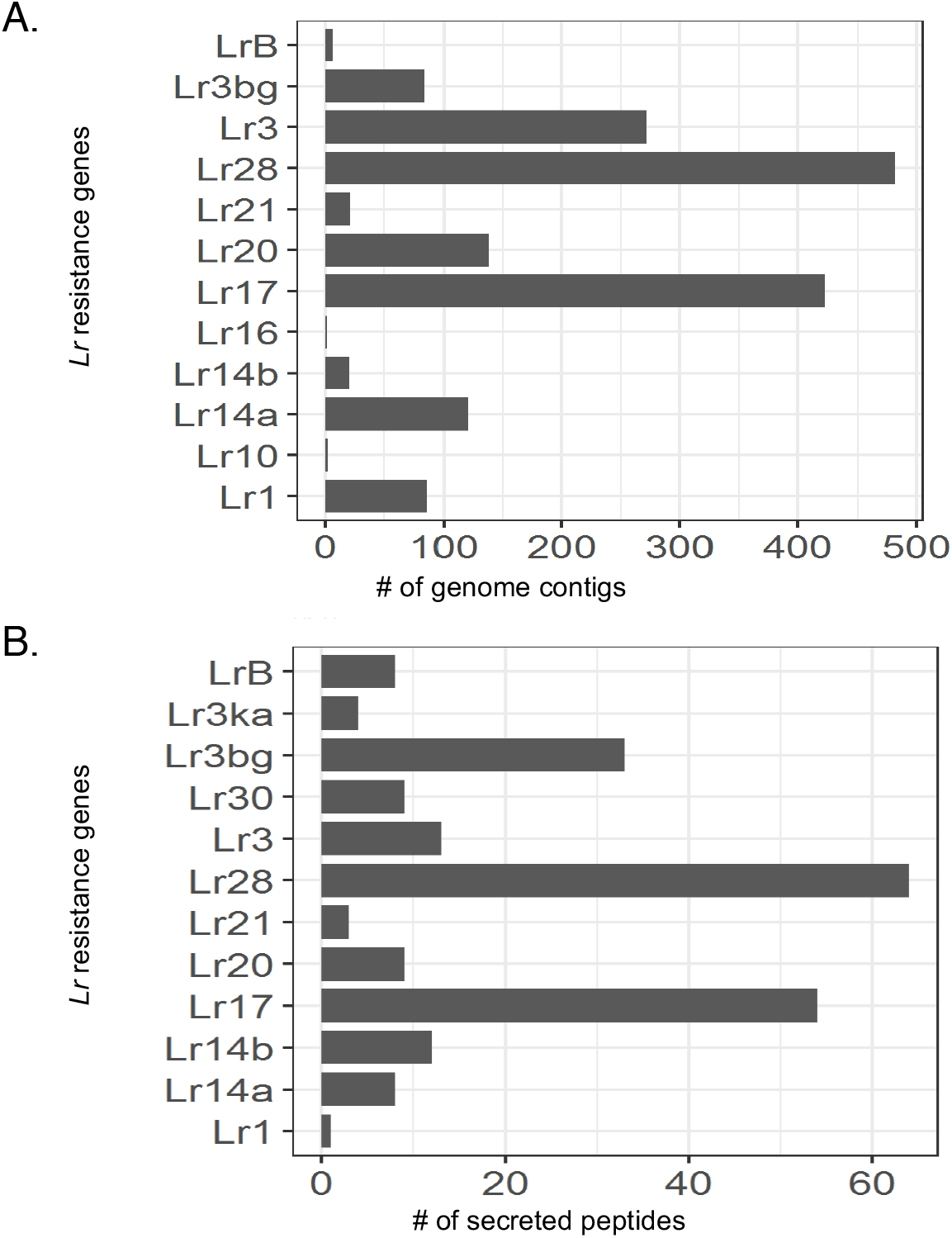
Genome-wide association analysis (GWAS) determining association of SNPs to changes in virulence to leaf rust resistance genes (Lr) A) Number of genome contigs that are associated with changes in virulence in all 121 isolates to individual resistance genes. B) Number of secreted peptides in all 121 isolates associated with changes in virulence. In both cases, the false discovery rate = 0.05.

## DISCUSSION

The lineage groups in North America are highly associated with avirulence/virulence to specific leaf rust resistance genes. In the absence of sexual recombination, clonal reproduction of urediniospores has maintained the association between molecular genotype and virulence. In a previous study based on allelic variation at 23 SSR loci (Ordonez and Kolmer, 2009), isolates with the characteristic avirulence/virulence as the isolates in this study were placed in the NA1, NA2, NA3, NA4, NA5 groups, and in a separate group of isolates that had high virulence to durum wheat. The NA1, NA2, NA4, and NA5 groups are long-standing in North America, as isolates with the characteristic virulence (Johnston et al 1968) can be found dating to the 1950s. Isolates in NA3 first become common in the mid 1990s after hard red winter wheat cultivars with *Lr37* were widely grown in the southern Great Plains (Ordonez and Kolmer, 2009). In the current *P. triticina* population in the U.S., isolates with the characteristic virulence of NA3 account for the large majority of all isolates (Kolmer, 2019). Two isolates were assigned to different NA groups based on SNP genotype compared to SSR genotypes. Isolate PBJSF_84VA was previously placed in the NA2 group based on SSR genotype, however the complete genome sequence data placed this isolate with isolate PBJL_PRTUS6 in the NA6 group. Similarly, isolate TCTDL_03VA was placed in NA1 based on SSR genotype, however based on complete genome sequence this isolate was not closely related to any other isolates that were sequenced in this study. Isolate Durum_09AZ103A was collected from durum wheat in Arizona in 2009. Surprisingly this isolate had an SNP genotype that placed it with isolates from Europe, indicating the high probability of this isolate being a recent introduction. The isolates from Europe that were sequenced in this study were distinct for SNP genotype from all of the other North American isolates. In a previous study, these same European isolates were also very distinct for SSR genotype compared to the North American isolates (Kolmer et al 2019).

In this study, all of the isolates were sequenced using short read technology. When BBBDB race 1-1 was re-sequenced and aligned back to the genome reference sequence, 273,385 SNPs were identified. The reference genome (Cuomo et al. 2017) was assembled using the Arachne (HybridAssemble; Jaffe et al., 2003) program, and it is likely that since the genome is dikaryotic significant regions sequenced from each nucleus were collapsed into one contig. The high number of SNPs from the alignment of BBBDB race 1-1 back to the reference genome is likely due to sequence from each genome being aligned to a contig region and the GATK program identifying differences caused by the two genomes. The high number of SNPs of the re-sequenced BBBDB_race 1-1 and the other isolates in the NA1 group relative to the reference genome is most probably caused by the assembly of the two genomes into a single genome. The high SNP heterozygosity of the groups from common wheat may also be a result of the genomes being collapsed into a single genome. Wu et al. (2017) re-sequenced 20 isolates of *P. triticina* from Australia that were aligned with the reference genome. The 20 isolates averaged 404,690 SNPs with 87% heterozygosity relative to the reference genome. Recently, the genome assembly of *P. graminis* f.sp. *tritici* was sorted into two groups representing the two nuclei. Using long reads, chromosome association, and specific algorithms, Feng et al. (2019) was able to sort contigs into their respective genomes. Kolmer et al., (2020) noted much lower heterozygosity on the order to 8-12% of SNPs in a large collection of *P. triticina* isolates that were characterized with the genotype-by-sequence (GBS) approach. The reads (100 bp, single end) of isolates in the GBS study were also aligned with the BBBDB race 1-1 reference genome, however the alignment, genotype data, VCF file, and haplotypes were extracted using different software programs. Differences in the mapping, SNP calling, threshold levels for heterozygotes, genome coverage, regions of the genome sequenced, depth of coverage, and assembly programs may also contribute the discrepancies in results between studies. Interestingly, both groups of isolates from durum wheat and the single isolate from *A. speltoides* had much lower levels of SNP heterozygosity compared to the isolates from common wheat.

Results of the whole genome sequence confirmed the genetic relationships between the isolate virulent to the diploid wheat progenitor *A. speltoides*, and forms of *P. triticina* specialized for virulence to tetraploid durum wheat and hexaploid common wheat. The maximum parsimony neighbor joining tree indicated that the *A. speltoides* isolate was most closely related to the ETH-Durum isolates, followed by the worldwide isolates virulent to durum wheat, and then isolates virulent to common wheat. In a coalescence analysis based on sequence at 15 loci, Liu et al 2014 determined that the form found on *A. speltoides* was the ancestral form of *P. triticina*, followed by the ETH-Durum isolates, and then by the isolates virulent to common wheat and durum wheat. Kolmer et al., (2020) using a molecular clock approach in the GBS study also found the same phylogenetic relationships between the different forms of *P. triticina*. The ETH-Durum isolates are highly virulent to durum wheat, but are avirulent to almost all common wheat cultivars, and have been found only in Ethiopia (Kolmer and Acevedo 2016). Ethiopia is a secondary center of origin of tetraploid wheat, where landrace cultivars of emmer and durum wheat are still grown (Eticha et al. 2006). The ETH-Durum isolates may be a remnant of the *P. triticina* population that existed before the widespread cultivation of hexaploid common wheat cultivars. The unique host environment of the landrace tetraploid wheats may allow the ETH-Durum isolates to survive in Ethiopia. The two isolates in NA-4 have virulence phenotypes that are found on the diploid wheat relative, *A. cylindrica* in the southern Great Plains of the U.S. The *P. triticina* population in Europe were previously determined to be highly diverse for SSR genotype groups (Kolmer et al 2012). Somatic recombination has been hypothesized as a source of variation in cereal rust fungi that reproduce clonally (Park et al 1999; Park and Wellings, 2012).

Host populations are important drivers of differentiation of plant pathogens at a species level and also for forms and genotypes within a species (Gladieux et al., 2017). The evolution of wheat from diploid progenitors to tetraploid and hexaploid cultivars was tracked by *P. triticina* (Liu et al 2014). Isolates of *P. triticina* that are virulent to hexaploid common wheat have many different virulence phenotypes that have been selected by leaf rust resistance genes (Kolmer 2019). Sequence variation in genes associated with pathogenicity and gene-for-gene interactions could possibly be used to differentiate pathogen genotypes for virulence to host resistance genes. However, even though isolates in the different NA groups differ for characteristic virulence to *Lr2a*, *Lr3, Lr3bg, Lr17, Lr28*, and *LrB,* there was no difference in the overall genetic relationship between isolates with groupings based on total genomic SNPs compared to groupings based on SNPs located in predicted secreted proteins associated with effector avirulence genes. Likewise, there was little difference in the nucleotide diversity, differentiation across contigs, and genetic similarity in relation to phenotypic similarity of total genomic SNPs and SNPs in secreted peptides. Contigs that had SNPs associated with virulence to *Lr17* and *Lr28* were the most numerous in both the total genomic SNPs and also the SNPs in the predicted secreted peptides. Since the current *P. triticina* population reproduces clonally, genomic SNPs associated with an NA group may not necessarily be linked with an effector gene, because mutations that occur throughout the genome are essentially linked, such as those found associated with *Lr17* virulence. Similarly, the SNPs in predicted secreted proteins may be widely distributed in the genome and not specific to effectors that interact with isolate specific *Lr* genes.

## CONCLUSIONS

In the absence of sexual recombination, mutations in noncoding and coding regions in *P. triticina* have accumulated, resulting in high levels of SNP and SSR (Kolmer et al 2019) heterozygosity, genetically differentiated groups, and a consistent association between virulence/avirulence and molecular genotypes. Despite the lack of sexual recombination with the resultant accumulation of mutations in coding regions different virulent phenotypes of *P. triticina* have continued to adapt to host resistance genes in both durum and common wheat cultivars and spread throughout the world. The forces of recurrent mutation, genetic drift, and selection by host resistance genes have together differentiated and diversified populations of *P. triticina* worldwide.

## METHODS

### Isolate phenotyping

Single uredinial isolates of *P. triticina* were increased on susceptible seedlings of *T. aestivum* variety ‘Little Club’ wheat (CI 4066; Clark et al, 1926), and then inoculated onto 7-day old seedlings of Thatcher near-isogenic differential lines that differ for single leaf rust resistance genes. Thatcher lines with genes *Lr1, Lr2a, Lr2c, Lr3, Lr9, Lr16, Lr24, Lr26, Lr3ka, Lr11, Lr17, Lr30, LrB, Lr10, Lr14a, Lr18, Lr3bg, Lr14b, Lr20*, *Lr21, Lr28* and *Lr39* were used to characterize the isolates for virulence. Protocols for rust inoculation, incubation, and collection of urediniospores, and greenhouse conditions were as previously described (Kolmer and Hughes 2018). The isolates were rated for virulence to the individual Thatcher differential lines 10-12 days after inoculation. Isolates with infection types 3-4 (moderate to large uredinia lacking necrosis or chlorosis) were considered virulent, and infection types 0-2^+^ (immune response with no sign of infection to moderate size uredinia with prominent chlorosis) were considered avirulent to the particular Thatcher line. Each isolate was given a 20-digit binary number based on the avirulent/virulent response to the 20 Thatcher differential lines.

### DNA Extraction

Seed of *Triticum aestivum* L. variety “Little Club” was planted in six 10” × 10” × 2” aluminum cake pans containing Metro Mix 360 soil media (SunGro). At the two-leaf stage, seedlings were inoculated with 25 mg of urediniospores from each isolate by suspending the spores in 3 ml of Soltrol 170 paraffin oil (Phillips Petroleum, Bartlesville, OK). Seedlings were fogged with an atomizer using 40 psi of compressed air and placed in a dew chamber (I-36DL, Percival Scientific, Perry, IA) overnight at 100% humidity and 20-22°C with lights off. Plants were placed back into a controlled environment chamber for 16 h/8 h day/night light cycle and 20 °C. After 11-14 d, spores were collected using a cyclonic Kramer-Collins spore collector (Tallgrass Solutions, Inc. Manhattan, KS). Urediniospores were germinated overnight by evenly scattering 0.5 g of spores over 300 ml of 1X germination solution contained in 20 × 30 cm Pyrex^®^ baking pan (Webb et al. 2006). DNA was isolated as described in Cuomo et al., (2017). Spore mats were skimmed off the surface of the germination solution, washed 3X with ddH_2_O, dried in a Buchner funnel, and stored at −80 °C. From each mat, 350 mg of tissue was ground using liquid N_2_. DNA was isolated using the OmniPrep™ DNA isolation Kit according to the recommended protocol for large tissue samples (Q-Biosciences, St. Louis MO, Cuomo et al., 2017).

### Isolate sequencing and variance analysis

DNA samples were sent to the Broad Institute (Cambridge, MA) for sequencing and analysis. From each sample, 200 bp insert Illumina (Illumina, San Diego, CA) libraries were constructed and ligated with a unique index barcode. All libraries were pooled and sequenced with eight lanes of HiSeq 2000, 101 base-pairedend reads. Illumina reads were aligned against version 2 of the *P. triticina* race 1-1 assembly (BBBDB; NCBI accession PRJNA36323) using BWA (Version: 0.5.9). Alignments were filtered requiring a mapping quality of 30 using SAMtools view (Li et al., 2009; Li and Durbin, 2010). Regions with high insertion/deletion counts, which could be misaligned, were identified using GATK (version v2.1.9; Makenna et al., 2010) RealignerTargetCreator and realigned using the GATK IndelRealigner. Single-nucleotide polymorphisms (SNPs) were identified using the GATK UnifiedGenotyper and filtered using GATK VariantFiltration (version 3) best practices for hard filters (QD < 2.0, MQ < 30.0, FS > 60.0, HS > [μ+2σ] MQRankSum < −12.5, ReadPosRankSum < −8.0), except that Haplotype Score (HS) was filtered above the mean HS plus two standard deviations. A single table of SNP calls in all strains was constructed, and at positions where no call was present in a strain the sequence data was re-evaluated to predict homozygous reference genotype calls. Using the pileup output from SAMtools, homozygous reference positions were called requiring a depth of at least eight and reference allele frequency >80%. Phylogenetic relationship based on heterozygous and homozygous SNP differences were inferred using PAUP Maximum Parsimony and ‘hetequal’ substitution model (Swafford 1998; 2002). DNA sequence data were deposited at NCBI BioSample PRJNA77809, and individual race SRA run numbers are listed in Supplemental Table 1. PLINK1.9 was used for the calculation of principal components, and identity-by-descent and identity-by-state (IBD/IBS). VCFtools (https://vcftools.github.io/) was used to calculate *F_ST_* and diversity nucleotide *π*. Genome-wide association (GWAS) was performed through compressed mixed linear model (MLM) in GAPIT with the first 3 principal components as population structure (Zhang et al., 2010). SNPs were considered a significant GWAS association at a false data rate of adjusted P-value < 0.05.

## Declarations

### Ethics Approval

NA

### Consent for publication

All authors have consented for submission for publication.

### Data Availability

All data has been deposited at NCBI under Bioproject PRJNA77809. Individual isolates SRA sequence deposits are listed in Supplementary Table 1.

### Competing interests

Authors declare no competing interests.

### Funding

This work was funded through USDA CSREE awards 2008-35600-04693 and 2009-65109-05916 and USDA-CRIS project 3020-21000-10-00D

### Authors’ Contributions

JPF, KM, and JAK Planned and developed experiments, isolated DNA, analyzed results and co-wrote the manuscript. SS and CAC planned and developed experiments, sequenced the samples, analyzed results and co-wrote the manuscript. GB provided data and co-wrote the manuscript.

## Acknowledgements

We thank the Broad Institute Genomics Platform for generating all sequence data for this study. The authors would like to thank Robert Bowden and Myron Bruce for their assistance in preparation of the manuscript and technical guidance.

Mention of trade names or commercial products in this publication is solely for the purpose of providing specific information and does not imply recommendation or endorsement by the U.S. Department of Agriculture. USDA is an equal opportunity provider and employer.

## Notes

### Competing Interest Statement

The authors have declared no competing interest.

