## Supplemental table 1 for "Whole-genome sequencing of multiple isolates of *Puccinia triticina* reveals asexual lineages evolving by recurrent mutations"

Supplemental Table 1. Summary of SNPs within 121 isolates of *Puccinia triticina* . Also included is the SRA run number, lineage, country of origin, number of SNPs withing specific regions of the genome, number of SNPs that are homozygous or heterozygous, virulences and references of previous data on the isolate.

| Isolate_Name-TreeCode | SRA Run number | Lineage | Country | %_of_Assembly_Covered_by_the_reads | Avg.Depth_of_Cover_(X) | SNPs | Intergenic | Intronic | p5UTR | p3UTR | Synonymous | Non-Synonymous | Read-Through | Nonsense | Hom | Het | Hom% | Het% | Virulence | Reference |
|---|---|---|---|---|---|---|---|---|---|---|---|---|---|---|---|---|---|---|---|---|
| Pt_ISR_850 | SRR11070886 | *Ae. speltoidies* | Israel | 78 | 33 | 996,187 | 780,886 | 49,700 | 11,594 | 22,589 | 60,101 | 69,440 | 372 | 1,505 | 645,298 | 350,889 | 65 | 35 | | |
| **Genome** | SRR403996 | **NA1** | USA | **99** | **48** | **273,526** | **229,385** | **8,450** | **1,953** | **3,527** | **11,198** | **18,395** | **127** | **491** | **4,156** | **269,370** | **2** | **98** | 14a, 14b, 20 | Ordonez and Kolmer 2009 |
| BBBD_04NE346_2 | SRR630042 | NA1 | USA | 99 | 51 | 284,163 | 238,499 | 8,774 | 2,010 | 3,629 | 11,511 | 19,092 | 136 | 512 | 4,447 | 279,716 | 2 | 98 | 14a, 14b, 20 | Ordonez and Kolmer 2009 |
| LBBKK_99NC | SRR630044 | NA1 | USA | 99 | 51 | 355,420 | 299,367 | 10,697 | 2,449 | 4,432 | 14,033 | 23,561 | 148 | 733 | 18,891 | 336,529 | 5 | 95 | 1, 10, 14a, 18, 14b, 20, 28 | Ordonez and Kolmer 2009 |
| PBDJ_84PA1 | SRR11080357 | NA1 | USA | 99 | 53 | 372,194 | 312,970 | 11,443 | 2,655 | 4,869 | 14,853 | 24,566 | 159 | 679 | 12,123 | 360,071 | 3 | 97 | 1, 2c, 3, 17, 10, 14a, 3bg, 20, 28 | Ordonez and Kolmer 2009 |
| FBMTK_04GA88_03 | SRR630059 | NA2 | USA | 98 | 46 | 420,866 | 351,885 | 13,152 | 3,176 | 5,781 | 17,489 | 28,400 | 172 | 811 | 56,173 | 364,693 | 13 | 87 | 2c, 3, 3ka, 30, B, 10, 14a, 18, 14b, 20, 28 | Ordonez and Kolmer 2009 |
| FLLLK_82TX2 | SRR630068 | NA2 | USA | 99 | 51 | 420,650 | 351,994 | 13,180 | 3,165 | 5,741 | 17,352 | 28,268 | 171 | 779 | 49,691 | 370,959 | 12 | 88 | 2c, 3, 9, 3ka, B, 14b, 20, 28 | Ordonez and Kolmer 2009 |
| NBBGT_11US182_1 | SRR11479797 | NA2 | USA | 98 | 42 | 440,318 | 368,962 | 13,750 | 3,191 | 5,936 | 18,002 | 29,459 | 185 | 833 | 40,556 | 399,762 | 9 | 91 | 1, 2c, B, 10, 14a, 18, 14b, 20, 28 | Kolmer et al 2013 |
| NBBR_98WA1 | SRR11080356 | NA2 | USA | 98 | 39 | 410,643 | 343,760 | 12,842 | 3,004 | 5,561 | 16,899 | 27,659 | 170 | 748 | 61,496 | 349,147 | 15 | 85 | 1, 2c, B, 10, 18, 14b, 20, 28 | Ordonez and Kolmer 2009 |
| NBBRG_11US203_2B | SRR11071604 | NA2 | USA | 97 | 42 | 436,140 | 365,543 | 13,534 | 3,150 | 5,913 | 17,789 | 29,210 | 175 | 826 | 61,307 | 374,833 | 14 | 86 | 1, 2c, B, 10, 18, 14b, 20, 28 | Kolmer et al 2013 |
| NBBSG_11US179_2A | SRR11074663 | NA2 | USA | 97 | 35 | 427,924 | 358,386 | 13,329 | 3,141 | 5,801 | 17,550 | 28,740 | 177 | 800 | 60,233 | 367,691 | 14 | 86 | 1, 2c, B, 10, 14a, 14b, 20, 28 | Kolmer et al 2013 |
| NBBTG_11US203_3B | SRR11071603 | NA2 | USA | 97 | 36 | 429,221 | 359,465 | 13,391 | 3,117 | 5,790 | 17,664 | 28,820 | 174 | 800 | 61,595 | 367,626 | 14 | 86 | 1, 2c, B, 10, 14a, 18, 14b, 20, 28 | Kolmer et al 2013 |
| NBGKK_82MN116_1 | SRR11080373 | NA2 | USA | 99 | 36 | 404,124 | 337,996 | 12,752 | 2,991 | 5,483 | 16,697 | 27,271 | 172 | 762 | 33,986 | 370,138 | 8 | 92 | 1, 2c, 11, 17, 30, 10, 14a, 18, 14b, 20, 28 | Ordonez and Kolmer 2009 |
| PBLR_91MI | SRR11080364 | NA2 | USA | 99 | 50 | 423,147 | 354,292 | 13,148 | 3,181 | 5,816 | 17,438 | 28,312 | 174 | 806 | 61,940 | 361,207 | 15 | 85 | 1, 2c, 3, 3ka, B, 10, 18, 14b, 20,28 | Ordonez and Kolmer 2009 |
| PBMGK_87NY626_2 | SRR11080363 | NA2 | USA | 99 | 42 | 410,299 | 343,084 | 12,815 | 3,104 | 5,657 | 16,970 | 27,711 | 167 | 791 | 46,246 | 364,053 | 11 | 89 | 1, 2c, 3, 3ka, 30, 10, 14b, 20, 28 | Ordonez and Kolmer 2009 |
| PNMR_94LA101 | SRR11080362 | NA2 | USA | 99 | 52 | 426,273 | 357,306 | 13,188 | 3,190 | 5,805 | 17,434 | 28,368 | 174 | 808 | 58,158 | 368,115 | 14 | 86 | 1, 2c, 3, 9, 24, 3ka, 30, B, 10, 18, 14b, 20, 28 | Ordonez and Kolmer 2009 |
| MBDS_06CAN32_1 | SRR630053 | NA3 | USA | 98 | 47 | 340,570 | 286,685 | 10,228 | 2,505 | 4,398 | 13,554 | 22,392 | 154 | 654 | 83,608 | 256,962 | 25 | 75 | 1, 3, 17, B, 10, 14a, 3bg, 14b, 20 | Ordonez and Kolmer 2009 |
| MBDSD_11US129_2 | SRR11071608 | NA3 | USA | 96 | 38 | 352,598 | 297,102 | 10,565 | 2,534 | 4,466 | 13,947 | 23,174 | 154 | 656 | 85,729 | 266,869 | 24 | 76 | 1, 3, 3ka, 17, 30, B, 10, 14a, 3bg, 14b, 20, 39 | Kolmer et al 2013 |
| MBDSD_11US144_2 | SRR11071607 | NA3 | USA | 96 | 37 | 351,411 | 295,986 | 10,558 | 2,529 | 4,473 | 13,913 | 23,135 | 154 | 663 | 82,977 | 268,434 | 24 | 76 | 1, 3, 3ka, 17, 30, B, 10, 14a, 3bg, 14b, 20, 39 | Kolmer et al 2013 |
| MBPSB_11US159_1 | SRR11071596 | NA3 | USA | 96 | 32 | 343,499 | 288,895 | 10,337 | 2,512 | 4,384 | 13,783 | 22,797 | 154 | 637 | 86,066 | 257,433 | 25 | 75 | 1, 3, 3ka, 17, 30, B, 10, 14a | Kolmer et al 2013 |
| MBTNB_11US148_2 | SRR11071592 | NA3 | USA | 96 | 30 | 344,027 | 289,615 | 10,297 | 2,519 | 4,391 | 13,673 | 22,717 | 153 | 662 | 79,588 | 264,439 | 23 | 77 | 1, 3, 3ka, 11, 17, 30, B,14a, 3bg, 14b, 20 | Kolmer et al 2013 |
| MBTNB_11US213_1 | SRR11071591 | NA3 | USA | 95 | 29 | 341,047 | 286,996 | 10,315 | 2,487 | 4,360 | 13,575 | 22,503 | 152 | 659 | 84,489 | 256,558 | 25 | 75 | 1, 3, 3ka, 11, 17, 30, B,14a, 3bg, 14b, 20 | Kolmer et al 2013 |
| MCDS_03CA360_2 | SRR630069 | NA3 | USA | 98 | 46 | 339,654 | 285,910 | 10,218 | 2,461 | 4,349 | 13,462 | 22,445 | 154 | 655 | 77,631 | 262,023 | 23 | 77 | 1, 3, 26, 17, B, 10, 14a, 3bg, 14b, 20 | Ordonez and Kolmer 2009 |
| MCDSB_11US185_2 | SRR11071590 | NA3 | USA | 96 | 31 | 346,275 | 291,168 | 10,521 | 2,558 | 4,421 | 13,882 | 22,913 | 153 | 659 | 82,659 | 263,616 | 24 | 76 | 1, 3, 26, 3ka, 17, 30, B, 10, 14a | Kolmer et al 2013 |
| MCDSD_11US034_2 | SRR11071589 | NA3 | USA | 96 | 47 | 358,888 | 302,789 | 10,616 | 2,601 | 4,549 | 14,068 | 23,416 | 155 | 694 | 86,685 | 272,203 | 24 | 76 | 1, 3, 26, 3ka, 17, 30, B, 10, 14a, 3bg, 14b, 20, 39 | Kolmer et al 2013 |
| MCJSS_04ND447 | SRR630045 | NA3 | USA | 98 | 51 | 342,119 | 288,050 | 10,239 | 2,490 | 4,382 | 13,553 | 22,581 | 148 | 676 | 84,571 | 257,548 | 25 | 75 | 1, 3, 26, 11, 17, B, 10, 14a, 3bg, 14b, 20 | Ordonez and Kolmer 2009 |
| MCPSB_11US230_1 | SRR11071588 | NA3 | USA | 96 | 46 | 357,000 | 301,177 | 10,609 | 2,570 | 4,502 | 14,027 | 23,291 | 159 | 665 | 85,891 | 271,109 | 24 | 76 | 1, 3, 26, 3ka, 17, 30, B, 10, 14a | Kolmer et al 2013 |
| MCPSS_04ND477 | SRR630063 | NA3 | USA | 98 | 51 | 342,974 | 288,677 | 10,356 | 2,489 | 4,393 | 13,569 | 22,651 | 160 | 679 | 79,781 | 263,193 | 23 | 77 | 1, 3, 26, 3ka, 17, 30, B, 10, 14a, 3bg, 14b, 20 | Ordonez and Kolmer 2009 |
| MCTNB_11US204_3 | SRR11071587 | NA3 | USA | 96 | 50 | 361,875 | 305,562 | 10,716 | 2,609 | 4,529 | 14,139 | 23,484 | 157 | 679 | 86,252 | 275,623 | 24 | 76 | 1, 3, 26, 3ka, 11, 17, 30, B,14a, 3bg, 14b, 20 | Kolmer et al 2013 |
| MCTNB_11US205_2 | SRR11071586 | NA3 | USA | 96 | 42 | 355,918 | 299,989 | 10,673 | 2,564 | 4,551 | 14,011 | 23,291 | 159 | 680 | 87,240 | 268,678 | 25 | 75 | 1, 3, 26, 3ka, 11, 17, 30, B,14a, 3bg, 14b, 20 | Kolmer et al 2013 |
| MCTSB_11US019_2 | SRR11080361 | NA3 | USA | 94 | 13 | 301,881 | 252,542 | 9,379 | 2,226 | 3,912 | 12,499 | 20,609 | 127 | 587 | 82,512 | 219,369 | 27 | 73 | 1, 3, 26, 3ka, 11, 17, 30, B, 10, 14a, 3bg, 14b, 20 | Kolmer et al 2013 |
| MCTSB_11US116_1 | SRR11479798 | NA5 | USA | 96 | 44 | 374,776 | 316,434 | 11,044 | 2,667 | 4,722 | 14,651 | 24,401 | 156 | 701 | 85,663 | 289,113 | 23 | 77 | 1, 3, 26, 3ka, 11, 17, 30, B, 10, 14a, 3bg, 14b, 20 | Kolmer et al 2013 |
| MDDSB_11US040_3 | SRR11071606 | NA3 | USA | 96 | 31 | 345,476 | 290,882 | 10,386 | 2,508 | 4,395 | 13,782 | 22,730 | 155 | 638 | 83,532 | 261,944 | 24 | 76 | 1, 3, 24, 17, B, 10, 14a, 3bg, 14b, 20 | Kolmer et al 2013 |
| MDFSB_11US187_2 | SRR11074672 | NA3 | USA | 96 | 30 | 343,398 | 289,225 | 10,234 | 2,505 | 4,383 | 13,634 | 22,616 | 147 | 654 | 85,209 | 258,189 | 25 | 75 | 1, 3, 24, 17, 30, B, 10, 14a, 3bg, 14b, 20 | Kolmer et al 2013 |
| MDPSB_11US122_1 | SRR11071605 | NA3 | USA | 93 | 10 | 267,919 | 224,056 | 8,240 | 1,978 | 3,521 | 11,183 | 18,277 | 131 | 533 | 78,579 | 189,340 | 29 | 71 | 1, 3, 24, 3ka, 17, 30, B, 10, 14a, 3bg, 14b, 20 | Kolmer et al 2013 |
| MFNSB_11US014_2 | SRR11080360 | NA3 | USA | 95 | 21 | 328,275 | 275,602 | 9,930 | 2,412 | 4,242 | 13,284 | 21,997 | 150 | 658 | 83,419 | 244,856 | 25 | 75 | 1, 3, 24, 26, 3ka, 17, B, 10, 14a, 3bg, 14b, 20 | Kolmer et al 2013 |
| MFNSB_11US029_3 | SRR11479796 | NA3 | USA | 96 | 46 | 357,858 | 301,757 | 10,698 | 2,577 | 4,535 | 14,021 | 23,419 | 163 | 688 | 82,012 | 275,846 | 23 | 77 | 1, 3, 24, 26, 3ka, 17, B, 10, 14a, 3bg, 14b, 20 | Ordonez and Kolmer 2009 |
| MFNSB_11US039_2 | SRR11080359 | NA3 | USA | 94 | 15 | 301,619 | 252,595 | 9,177 | 2,242 | 3,905 | 12,427 | 20,535 | 138 | 600 | 81,150 | 220,469 | 27 | 73 | 1, 3, 24, 26, 3ka, 17, B, 10, 14a, 3bg, 14b, 20 | Kolmer et al 2013 |
| MFNSB_11US074_2 | SRR11080358 | NA3 | USA | 96 | 30 | 344,459 | 289,623 | 10,384 | 2,525 | 4,406 | 13,744 | 22,949 | 154 | 674 | 84,959 | 259,500 | 25 | 75 | 1, 3, 24, 26, 3ka, 17, B, 10, 14a, 3bg, 14b, 20 | Kolmer et al 2013 |
| MFNSB_11US220_3 | SRR11074665 | NA3 | USA | 97 | 95 | 377,477 | 319,312 | 11,170 | 2,688 | 4,692 | 14,453 | 24,282 | 160 | 720 | 85,307 | 292,170 | 23 | 77 | 1, 3, 24, 26, 3ka, 17, B, 10, 14a, 3bg, 14b, 20 | Kolmer et al 2013 |
| MFPS_06MN268 | SRR630058 | NA3 | USA | 99 | 49 | 340,412 | 286,608 | 10,226 | 2,473 | 4,386 | 13,481 | 22,408 | 153 | 677 | 65,285 | 275,127 | 19 | 81 | 1, 3, 24, 26, 3ka, 17, 30, B, 10, 14a | Ordonez and Kolmer 2009 |
| MHDS_03OH237 | SRR630064 | NA3 | USA | 98 | 45 | 337,866 | 284,242 | 10,164 | 2,476 | 4,374 | 13,482 | 22,317 | 153 | 658 | 83,822 | 254,044 | 25 | 75 | 1, 3, 16, 26, 17, B, 10, 14a, 3bg, 14b, 20 | Ordonez and Kolmer 2009 |
| MLDS_06MN384 | SRR630072 | NA3 | USA | 98 | 47 | 341,664 | 287,669 | 10,294 | 2,503 | 4,374 | 13,567 | 22,449 | 159 | 649 | 80,839 | 260,825 | 24 | 76 | 1, 3, 9, 17, B, 10, 14a, 3bg, 14b, 20,39 | Ordonez and Kolmer 2009 |
| MLDSD_11US025_3 | SRR11074664 | NA3 | USA | 96 | 36 | 350,469 | 295,259 | 10,486 | 2,524 | 4,452 | 13,953 | 22,967 | 152 | 676 | 86,114 | 264,355 | 25 | 75 | 1, 3, 9, 17, B, 10, 14a, 3bg, 14b, 20, 39 | Kolmer et al 2013 |
| MLDSS_10MN1_2 | SRR630051 | NA3 | USA | 98 | 48 | 341,747 | 287,751 | 10,299 | 2,506 | 4,375 | 13,560 | 22,447 | 159 | 650 | 73,260 | 268,487 | 21 | 79 | 1, 3, 9, 17, B, 10, 14a, 3bg, 14b, 20,39 | Kolmer et al 2012 |
| MMDSD_11US075_1 | SRR11080355 | NA3 | USA | 96 | 38 | 354,787 | 299,238 | 10,608 | 2,559 | 4,468 | 13,967 | 23,131 | 160 | 656 | 86,843 | 267,944 | 24 | 76 | 1, 3, 9, 26, 17, B, 10, 14a, 3bg, 14b, 20, 39 | Kolmer et al 2013 |
| TBDS_04TX67 | SRR11080354 | NA3 | USA | 98 | 46 | 341,327 | 287,445 | 10,231 | 2,487 | 4,385 | 13,581 | 22,373 | 150 | 675 | 83,226 | 258,101 | 24 | 76 | 1, 2a, 2c, 3, 17, B, 10, 14a, 3bg, 14, 20 | Ordonez and Kolmer 2009 |
| TCJSB_11US078_1 | SRR11071599 | NA3 | USA | 96 | 32 | 347,795 | 292,649 | 10,389 | 2,530 | 4,435 | 14,131 | 22,846 | 149 | 666 | 85,843 | 261,952 | 25 | 75 | 1,2a,2c,3, 26, 3ka, 17, B, 10, 14a, 3bg, 14b, 20 | Kolmer et al 2013 |
| TCPSS_04KS213 | SRR11071598 | NA3 | USA | 99 | 49 | 340,468 | 286,678 | 10,251 | 2,464 | 4,406 | 13,469 | 22,391 | 151 | 658 | 78,569 | 261,899 | 23 | 77 | 1, 2a, 2c, 3, 26, 3ka, 17, 30, B, 10, 14a, 3bg, 14b, 20 | Ordonez and Kolmer 2009 |
| TCTSB_11US189_3 | SRR11074662 | NA3 | USA | 95 | 27 | 338,198 | 284,835 | 10,094 | 2,417 | 4,321 | 13,454 | 22,311 | 155 | 611 | 84,204 | 253,994 | 25 | 75 | 1,2a,2c,3, 26, 3ka, 11, 17, 30, B, 10, 14a, | Kolmer et al 2013 |
| TDFSS_04TX86 | SRR11080381 | NA3 | USA | 98 | 53 | 344,421 | 290,209 | 10,340 | 2,514 | 4,413 | 13,578 | 22,551 | 151 | 665 | 86,315 | 258,106 | 25 | 75 | 1, 2a, 2c, 3, 24, 17, 30, B, 10, 14a, 3bg, 14, 20 | Ordonez and Kolmer 2009 |
| SBDB_04MN364 | SRR11080380 | NA4 | USA | 97 | 58 | 315,088 | 266,580 | 9,012 | 2,085 | 3,726 | 12,153 | 20,739 | 130 | 663 | 106,394 | 208,694 | 34 | 66 | 1, 2a, 2c, 17,14b | Ordonez and Kolmer 2009 |
| SBDGD_10MT13_1 | SRR11080379 | NA4 | USA | 97 | 45 | 304,977 | 257,846 | 8,717 | 2,066 | 3,629 | 11,847 | 20,113 | 129 | 630 | 103,106 | 201,871 | 34 | 66 | 1, 2a, 2c, 17, 10, 14b | Ordonez and Kolmer 2009 |
| KBBJK_84MN526_2 | SRR630060 | NA5 | USA | 99 | 49 | 417,004 | 349,053 | 13,083 | 3,107 | 5,651 | 17,156 | 28,013 | 172 | 769 | 20,955 | 396,049 | 5 | 95 | 2a, 2c, 3, 10, 14a, 14b, 20, 28 | Ordonez and Kolmer 2009 |
| KFBJG_97CA64_1 | SRR11074673 | NA5 | USA | 97 | 37 | 320,787 | 269,787 | 9,617 | 2,290 | 4,030 | 12,944 | 21,362 | 141 | 616 | 34,479 | 286,308 | 11 | 89 | 2a, 2c, 3, 24, 26, 10, 14a, 28, 14b, 20 | Ordonez and Kolmer 2009 |
| MBBJT_RACE5 | SRR630071 | NA5 | USA | 99 | 45 | 306,074 | 257,130 | 9,297 | 2,176 | 3,867 | 12,442 | 20,423 | 139 | 600 | 30,885 | 275,189 | 10 | 90 | 1, 3, 10, 14a, 3bg, 14b, 20, 28 | Ordonez and Kolmer 2009 |
| MBGDK_99NC23_1B | SRR11479799 | NA5 | USA | 99 | 46 | 312,142 | 262,422 | 9,390 | 2,209 | 3,947 | 12,576 | 20,848 | 142 | 608 | 28,747 | 283,395 | 9 | 91 | 1, 3, 11, 14a, 14b, 20, 28 | Ordonez and Kolmer 2009 |
| MBRL_KSUSDA1 | SRR630048 | NA5 | USA | 99 | 46 | 313,143 | 263,651 | 9,318 | 2,186 | 3,913 | 12,518 | 20,789 | 141 | 627 | 34,492 | 278,651 | 11 | 89 | 1, 3, 3ka, 11, 30, B | |
| MJBJ_97NE406 | SRR630041 | NA5 | USA | 99 | 40 | 304,976 | 256,138 | 9,253 | 2,192 | 3,885 | 12,356 | 20,395 | 137 | 620 | 33,928 | 271,048 | 11 | 89 | 1, 3, 16, 24, 10, 14a, 14b, 20, 28 | Ordonez and Kolmer 2009 |
| PRTUS1_PRTUS1 | SRR11080378 | NA5 | USA | 99 | 57 | 314,880 | 264,929 | 9,496 | 2,210 | 3,947 | 12,647 | 20,886 | 143 | 622 | 29,016 | 285,864 | 9 | 91 | --- | |
| TBBGJ_11US042_1 | SRR11071602 | NA5 | USA | 98 | 37 | 345,705 | 291,050 | 10,378 | 2,477 | 4,371 | 13,836 | 22,773 | 154 | 666 | 26,917 | 318,788 | 8 | 92 | 1, 2a, 2c, 3, 10, 28,39, 14b | Kolmer et al 2013 |
| TBBGJ_11US136_1 | SRR11071601 | NA5 | USA | 98 | 50 | 336,190 | 283,656 | 9,977 | 2,318 | 4,151 | 13,299 | 21,999 | 145 | 645 | 32,591 | 303,599 | 10 | 90 | 1, 2a, 2c, 3, 10, 28, 39, 14b | Ordonez and Kolmer 2009 |
| TCBJG_11US053_3 | SRR11071600 | NA5 | USA | 97 | 30 | 314,404 | 264,204 | 9,478 | 2,220 | 3,957 | 12,699 | 21,068 | 142 | 636 | 34,031 | 280,373 | 11 | 89 | 1, 2a, 2c, 3, 26, 10, 14a, 28, 14b, 20 | Kolmer et al 2013 |
| TCBJK_04TX104 | SRR11080377 | NA5 | USA | 99 | 45 | 311,027 | 261,385 | 9,350 | 2,198 | 3,937 | 12,584 | 20,786 | 146 | 641 | 33,067 | 277,960 | 11 | 89 | 1, 2a, 2c, 3, 26, 10, 14a, 14b, 20, 28 | Ordonez and Kolmer 2009 |
| TCGJG_11US256_3 | SRR11071595 | NA5 | USA | 97 | 34 | 321,337 | 270,090 | 9,571 | 2,280 | 3,995 | 13,174 | 21,455 | 142 | 630 | 35,886 | 285,451 | 11 | 89 | 1, 2a, 2c, 3, 26, 11, 10, 14a, 28, 14b, 20 | Kolmer et al 2013 |
| TCRKG_11US057_1 | SRR11071597 | NA5 | USA | 97 | 41 | 325,842 | 274,448 | 9,722 | 2,280 | 4,048 | 12,982 | 21,592 | 144 | 626 | 38,026 | 287,816 | 12 | 88 | 1, 2a, 2c, 3, 26, 3ka, 11, 30, 10, 14a, 18, 28, 14b, 20 | Kolmer et al 2013 |
| TCRKG_11US116_2 | SRR11071594 | NA5 | USA | 97 | 38 | 323,017 | 271,945 | 9,609 | 2,257 | 4,006 | 12,843 | 21,593 | 141 | 623 | 36,079 | 286,938 | 11 | 89 | 1, 2a, 2c, 3, 26, 3ka, 11, 30, 10, 14a, 18, 28, 14b, 20 | Kolmer et al 2013 |

| | | | | | | | | | | | | | | | | | | | | |
|---|---|---|---|---|---|---|---|---|---|---|---|---|---|---|---|---|---|---|---|---|
| TDBG_08TX210 | SRR11080376 | NA5 | USA | 99 | 44 | 311,859 | 262,284 | 9,350 | 2,199 | 3,928 | 12,530 | 20,806 | 141 | 621 | 34,346 | 277,513 | 11 | 89 | 1, 2a, 2c, 3, 24, 10, 14b, 28 | Ordonez and Kolmer 2009 |
| TDBJG_11US154_1 | SRR11080375 | NA5 | USA | 98 | 32 | 339,459 | 285,230 | 10,326 | 2,447 | 4,274 | 13,702 | 22,674 | 154 | 652 | 26,340 | 313,119 | 8 | 92 | 1, 2a, 2c, 3, 24, 10, 14a, 14b, 20, 21, 28 | Kolmer et al 2013 |
| TDBJQ_11US026_1 | SRR11074661 | NA5 | USA | 97 | 33 | 319,260 | 268,456 | 9,526 | 2,283 | 3,995 | 12,860 | 21,356 | 145 | 639 | 30,882 | 288,378 | 10 | 90 | 1, 2a, 2c, 3, 24, 10, 14a, 21, 28, 14b, 20 | Kolmer et al 2013 |
| TDBJQ_11US091_2 | SRR11074660 | NA5 | USA | 97 | 38 | 323,599 | 272,550 | 9,620 | 2,241 | 4,029 | 12,910 | 21,475 | 143 | 631 | 35,582 | 288,017 | 11 | 89 | 1, 2a, 2c, 3, 24, 10, 14a, 21, 28, 14b, 20 | Kolmer et al 2013 |
| TFBGG_11US176_2 | SRR11074659 | NA5 | USA | 97 | 37 | 323,369 | 272,028 | 9,695 | 2,272 | 4,031 | 12,994 | 21,556 | 140 | 653 | 34,483 | 288,886 | 11 | 89 | 1, 2a, 2c, 3, 10, 28, 14b | Kolmer et al 2013 |
| TFBGQ_11US212_1 | SRR11071593 | NA5 | USA | 98 | 39 | 353,556 | 297,364 | 10,708 | 2,466 | 4,489 | 14,190 | 23,523 | 153 | 663 | 26,566 | 326,990 | 8 | 92 | 1, 2a, 2c, 3, 24, 26, 10, 14b, 21, 28 | Kolmer et al 2013 |
| TFBJQ_10MN3_1_2 | SRR11080374 | NA5 | USA | 99 | 35 | 303,935 | 254,757 | 9,234 | 2,179 | 3,878 | 12,657 | 20,504 | 143 | 583 | 27,110 | 276,825 | 9 | 91 | 1, 2a, 2c, 3, 24, 26, 10, 14b, 21, 20, 28 | Kolmer et al 2012 |
| THBJ_99ND588 | SRR11080372 | NA5 | USA | 99 | 55 | 314,384 | 264,229 | 9,520 | 2,210 | 3,963 | 12,653 | 21,041 | 143 | 625 | 34,872 | 279,512 | 11 | 89 | 1, 2a, 2c, 3, 16, 26, 10, 14a, 14b, 20, 28 | Ordonez and Kolmer 2009 |
| TJBGH_04OH300 | SRR11080371 | NA5 | USA | 99 | 50 | 326,337 | 274,314 | 9,822 | 2,337 | 4,197 | 13,098 | 21,787 | 151 | 631 | 29,142 | 297,195 | 9 | 91 | 1, 2a, 2c, 3, 16, 24, 10, 14b, 28 | Ordonez and Kolmer 2009 |
| TNBGJ_11US022_3 | SRR11074658 | NA5 | USA | 98 | 36 | 325,962 | 274,364 | 9,732 | 2,280 | 4,086 | 13,086 | 21,637 | 143 | 634 | 25,199 | 300,763 | 8 | 92 | 1, 2a, 2c, 3, 9, 24, 10, 28, 39, 14b | Kolmer et al 2013 |
| TNBGJ_11US045_2 | SRR11080370 | NA5 | USA | 94 | 9 | 221,165 | 184,543 | 6,881 | 1,595 | 2,803 | 9,302 | 15,495 | 110 | 436 | 37,412 | 183,753 | 17 | 83 | 1, 2a, 2c, 3, 9, 24, 10, 28, 39, 14b, 20 | Kolmer et al 2013 |
| TNBGJ_11US048_2 | SRR11074671 | NA5 | USA | 97 | 35 | 324,943 | 273,928 | 9,621 | 2,262 | 4,042 | 12,904 | 21,413 | 142 | 631 | 35,563 | 289,380 | 11 | 89 | 1, 2a, 2c, 3, 9, 24, 10, 28, 39, 14b | Kolmer et al 2013 |
| TNBGJ_11US130_2 | SRR11074670 | NA5 | USA | 98 | 28 | 331,495 | 278,410 | 10,063 | 2,368 | 4,136 | 13,419 | 22,310 | 151 | 638 | 25,472 | 306,023 | 8 | 92 | 1, 2a, 2c, 3, 9, 24, 10, 28, 39, 14b | Kolmer et al 2013 |
| TNBJJ_11US030_1 | SRR11074669 | NA5 | USA | 97 | 42 | 326,845 | 275,246 | 9,744 | 2,276 | 4,074 | 13,052 | 21,672 | 150 | 631 | 35,998 | 290,847 | 11 | 89 | 1, 2a, 2c, 3, 9, 24, 10, 14a, 28, 39, 14b, 20 | Kolmer et al 2013 |
| TNRJ_05TX27 | SRR11080369 | NA5 | USA | 99 | 41 | 306,885 | 258,025 | 9,260 | 2,146 | 3,876 | 12,348 | 20,473 | 143 | 614 | 35,542 | 271,343 | 12 | 88 | 1, 2a, 2c, 3, 9, 24, 3ka, 17, 30, 10, 14a, 28, 39, 14b, 20, 28 | Ordonez and Kolmer 2009 |
| TNRJJ_11US043_3 | SRR11074668 | NA5 | USA | 97 | 33 | 315,995 | 265,664 | 9,536 | 2,236 | 3,958 | 12,724 | 21,129 | 139 | 609 | 36,339 | 279,656 | 11 | 89 | 1, 2a, 2c, 3, 9, 24, 3ka, 17, 30, 10, 14a, 28, 39, 14b, 20, 28 | Kolmer et al 2013 |
| TNRJJ_11US098_2 | SRR11074667 | NA5 | USA | 97 | 35 | 321,066 | 270,204 | 9,585 | 2,262 | 3,920 | 12,905 | 21,412 | 147 | 631 | 34,366 | 286,700 | 11 | 89 | 1, 2a, 2c, 3, 9, 24, 3ka, 17, 30, 10, 14a, 28, 39, 14b, 20, 28 | Kolmer et al 2013 |
| TPBGJ_11US044_1 | SRR11074666 | NA5 | USA | 97 | 34 | 347,697 | 292,470 | 10,496 | 2,513 | 4,383 | 13,961 | 23,059 | 159 | 656 | 31,629 | 316,068 | 9 | 91 | 1, 2a, 2c, 3, 9, 24, 26, 10, 28, 39, 14b, 28 | Kolmer et al 2013 |
| TCTDL_03VA190 | SRR11080368 | --- | USA | 99 | 45 | 338,950 | 286,802 | 9,776 | 2,340 | 4,073 | 13,012 | 22,114 | 145 | 688 | 51,125 | 287,825 | 15 | 85 | 1, 2a, 2c, 3, 26, 3ka, 11, 17, 30, 14a, 3bg | Ordonez and Kolmer 2009 |
| PBJL_PRTUS6 | SRR11080367 | NA6 | USA | 98 | 48 | 347,829 | 292,096 | 10,601 | 2,538 | 4,516 | 14,034 | 23,227 | 160 | 657 | 71,422 | 276,407 | 21 | 79 | 1, 2c, 3, 11, 17, 14a, 18, 21 | This report |
| PBJSF_84VA | SRR11479801 | NA2 | USA | 98 | 51 | 353,384 | 296,960 | 10,664 | 2,522 | 4,552 | 14,248 | 23,580 | 162 | 696 | 72,200 | 281,184 | 20 | 80 | 1, 2c, 3, 11, 17, 14a, 18, 20, 28 | Ordonez and Kolmer 2009 |
| DURUM_CA_1.2 | SRR630070 | DURUM | USA | 98 | 51 | 414,766 | 348,762 | 12,625 | 3,046 | 5,796 | 16,512 | 27,109 | 169 | 747 | 122,685 | 292,081 | 30 | 70 | B, 10, 14b, 20, 39 | Ordonez and Kolmer 2007 |
| DURUM_MX_14_3 | SRR630061 | DURUM | Mexico | 96 | 54 | 415,245 | 349,342 | 12,653 | 3,062 | 5,808 | 16,438 | 27,049 | 165 | 728 | 204,904 | 210,341 | 49 | 51 | B, 10, 14b, 20, 39 | Ordonez and Kolmer 2007 |
| FCBQB_CHL08_5_3 | SRR630062 | DURUM | Chile | 98 | 47 | 414,060 | 348,153 | 12,615 | 3,075 | 5,794 | 16,427 | 27,050 | 179 | 767 | 137,600 | 276,460 | 33 | 67 | 2c, 3, 26, 3ka, 11 | Ordonez and Kolmer 2007 |
| Pt_ARG_12.1 | SRR11070895 | DURUM | Argentina | 93 | 28 | 406,099 | 340,316 | 12,640 | 3,063 | 5,769 | 16,434 | 26,996 | 165 | 716 | 200,122 | 205,977 | 49 | 51 | B, 10, 14b, 20, 39 | Ordonez and Kolmer 2007 |
| Pt_ARG_9.3 | SRR11070894 | DURUM | Argentina | 94 | 72 | 442,480 | 372,647 | 13,506 | 3,300 | 6,098 | 17,300 | 28,663 | 182 | 784 | 208,399 | 234,081 | 47 | 53 | B, 10, 14b, 20, 39 | Ordonez and Kolmer 2007 |
| Pt_ESP_12 | SRR11070885 | DURUM | Spain | 92 | 25 | 403,909 | 338,070 | 12,479 | 3,098 | 5,610 | 16,493 | 27,265 | 186 | 708 | 206,202 | 197,707 | 51 | 49 | 10, 14b, 20, 39 | Ordonez and Kolmer 2007 |
| Pt_ESP_27 | SRR11070884 | DURUM | Spain | 92 | 40 | 424,628 | 356,436 | 12,940 | 3,211 | 5,843 | 17,017 | 28,237 | 196 | 748 | 211,750 | 212,878 | 50 | 50 | 10, 14b, 20, 39 | Ordonez and Kolmer 2007 |
| Pt_ESP_30 | SRR11070883 | DURUM | Spain | 94 | 39 | 422,488 | 355,115 | 12,903 | 3,174 | 5,862 | 16,873 | 27,645 | 166 | 750 | 197,026 | 225,462 | 47 | 53 | B, 10, 14b, 20, 39 | Ordonez and Kolmer 2007 |
| Pt_ESP_4 | SRR11070882 | DURUM | Spain | 93 | 39 | 425,622 | 358,048 | 12,914 | 3,137 | 5,913 | 16,914 | 27,776 | 176 | 744 | 192,354 | 233,268 | 45 | 55 | B, 10, 14b, 20 | Ordonez and Kolmer 2007 |
| Pt_ETH_11D5_1 | SRR11070881 | DURUM | Ethiopia | 93 | 33 | 413,547 | 347,063 | 12,696 | 3,129 | 5,823 | 16,628 | 27,304 | 168 | 736 | 201,669 | 211,878 | 49 | 51 | B, 10, 14b, 20, 39 | Ordonez and Kolmer 2007 |
| Pt_FRA_4.3 | SRR11070889 | DURUM | France | 92 | 27 | 403,070 | 337,675 | 12,466 | 3,049 | 5,749 | 16,394 | 26,870 | 163 | 704 | 199,464 | 203,606 | 49 | 51 | B, 10, 14b, 20, 39 | Ordonez and Kolmer 2007 |
| Pt_FRA_5.3 | SRR11070888 | DURUM | France | 95 | 24 | 401,792 | 336,605 | 12,449 | 3,044 | 5,721 | 16,332 | 26,762 | 168 | 711 | 120,098 | 281,694 | 30 | 70 | B, 10, 14b, 20, 39 | Ordonez and Kolmer 2007 |
| Pt_ISR_1508 | SRR11070887 | DURUM | Israel | 93 | 27 | 428,243 | 359,218 | 13,201 | 3,262 | 5,968 | 17,289 | 28,370 | 173 | 762 | 170,433 | 257,810 | 40 | 60 | B, 10, 14b, 20, 39 | Ordonez and Kolmer 2007 |
| Pt_ETH_11D9_3 | SRR11070880 | ETH-DURUM | Ethiopia | 93 | 33 | 441,196 | 370,961 | 13,203 | 3,222 | 5,999 | 17,671 | 29,169 | 178 | 793 | 203,319 | 237,877 | 46 | 54 | --- | Kolmer and Acevedo 2016 |
| Pt_ETH_125_2 | SRR11070879 | ETH-DURUM | Ethiopia | 93 | 36 | 446,946 | 376,335 | 13,251 | 3,243 | 6,061 | 17,723 | 29,354 | 180 | 799 | 204,520 | 242,426 | 46 | 54 | --- | Kolmer and Acevedo 2016 |
| Pt_ETH_16.1 | SRR11070878 | ETH-DURUM | Ethiopia | 93 | 32 | 411,203 | 346,516 | 12,194 | 2,988 | 5,438 | 16,242 | 26,979 | 172 | 674 | 221,328 | 189,875 | 54 | 46 | --- | Ordonez and Kolmer 2007 |
| Pt_ETH_4.1 | SRR11070893 | ETH-DURUM | Ethiopia | 96 | 33 | 476,566 | 399,670 | 14,416 | 3,565 | 6,615 | 19,262 | 31,992 | 202 | 844 | 136,340 | 340,226 | 29 | 71 | --- | Ordonez and Kolmer 2007 |
| Pt_ETH_4090_3 | SRR11070892 | ETH-DURUM | Ethiopia | 94 | 34 | 474,770 | 398,459 | 14,304 | 3,542 | 6,555 | 19,154 | 31,694 | 195 | 867 | 169,405 | 305,365 | 36 | 64 | --- | Ordonez and Kolmer 2007 |
| Pt_ETH_4114_7 | SRR11070891 | ETH-DURUM | Ethiopia | 91 | 38 | 423,210 | 356,595 | 12,505 | 3,127 | 5,600 | 16,721 | 27,781 | 175 | 706 | 260,292 | 162,918 | 62 | 38 | --- | Ordonez and Kolmer 2007 |
| Pt_ETH_6.1 | SRR11070890 | ETH-DURUM | Ethiopia | 94 | 37 | 479,917 | 403,054 | 14,367 | 3,636 | 6,573 | 19,297 | 31,934 | 200 | 856 | 169,390 | 310,527 | 35 | 65 | --- | Ordonez and Kolmer 2007 |
| SCPBJ_ISR173B | SRR11080366 | ME2 | Israel | 97 | 56 | 358,093 | 301,735 | 10,776 | 2,529 | 4,530 | 14,089 | 23,555 | 163 | 716 | 117,760 | 240,333 | 33 | 67 | 1, 2a, 2c, 26, 3ka, 17, 30, 14b, 20 | Kolmer et al 2011 |
| BBBQB_CZ20_09 | SRR630065 | EU1 | Czech-Slovakia | 98 | 47 | 439,466 | 368,177 | 13,752 | 3,288 | 5,827 | 17,888 | 29,509 | 192 | 833 | 91,856 | 347,610 | 21 | 79 | B, 10 | Kolmer et al 2011 |
| DGGQ_09GBR10_1 | SRR630050 | EU1 | Great Britain | 98 | 55 | 447,881 | 375,651 | 13,882 | 3,328 | 5,884 | 18,387 | 29,736 | 197 | 816 | 93,532 | 354,349 | 21 | 79 | 2c, 3, 16, 11, B, 10, 28 | Kolmer et al 2013 |
| DHJJ_09GB14.1 | SRR630057 | EU1 | Great Britain | 97 | 16 | 346,059 | 288,328 | 11,176 | 2,643 | 4,717 | 14,607 | 23,780 | 155 | 653 | 87,662 | 258,397 | 25 | 75 | 2c, 16, 26, 11, 17, B, 10, 14a, 28 | Kolmer et al 2013 |
| FCPSQ_09TUR11_1 | SRR630049 | EU2 | Turkey | 99 | 48 | 449,493 | 376,788 | 13,947 | 6,020 | 3,343 | 18,673 | 29,730 | 182 | 810 | 43,170 | 406,323 | 10 | 90 | 2c, 3, 26, 3ka, 11, 17, 30, B, 10, 14a, 3bg, 14b | Kolmer et al 2013 |
| FCPSQ_09TUR23_1 | SRR11479800 | EU2 | Turkey | 99 | 53 | 430,581 | 361,513 | 13,373 | 3,152 | 5,555 | 17,212 | 28,713 | 193 | 870 | 58,773 | 371,808 | 14 | 86 | 1, 2c, 3, 16, 3ka, 11, 17, 30, B, 10, 14a, 3bg, 14b, 20 | Kolmer et al 2013 |
| FCPNS_CZ18_09 | SRR630054 | EU4 | Czech-Slovakia | 97 | 21 | 345,141 | 288,302 | 10,847 | 2,478 | 4,519 | 14,517 | 23,644 | 157 | 677 | 93,773 | 251,368 | 27 | 73 | 2c, 3, 26, 3ka, 11, 17, 30, B, 14a, 3bg, 14b, 20 | Kolmer et al 2013 |
| FCPQQ_95SK2_2 | SRR630056 | EU4 | Czech-Slovakia | 98 | 55 | 401,650 | 337,353 | 12,338 | 2,854 | 5,199 | 16,126 | 26,784 | 176 | 820 | 101,384 | 300,266 | 25 | 75 | 2c, 3, 26, 3ka, 11, 17, 30, B, 10, 3bg, 14b | Kolmer et al 2013 |
| FCPSS_F95 | SRR630066 | EU4 | France | 98 | 56 | 403,382 | 338,999 | 12,330 | 2,864 | 5,213 | 16,183 | 26,800 | 187 | 806 | 102,116 | 301,266 | 25 | 75 | 2c, 3, 26, 3ka, 11, 17, 30, B, 10, 14a, 3bg, 14b, 20 | Kolmer et al 2013 |
| FHMQQ_CZ10_09 | SRR630055 | EU5 | Czech-Slovakia | 99 | 45 | 381,299 | 319,483 | 11,899 | 2,760 | 4,918 | 15,738 | 25,577 | 168 | 756 | 44,492 | 336,807 | 12 | 88 | 2c, 3, 16, 26, 3ka, 30, B, 10, 3bg, 14b | Kolmer et al 2013 |
| KCMQ_09TUR19_2_09 | SRR630067 | EU5 | Turkey | 99 | 52 | 432,017 | 361,785 | 13,497 | 3,236 | 5,805 | 17,695 | 29,032 | 181 | 786 | 50,501 | 381,516 | 12 | 88 | 2a, 2c, 3, 26, 3ka, 11, 3bg, 14b | Kolmer et al 2011 |
| FBPSQ_FR56 | SRR630046 | EU7 | France | 99 | 53 | 449,681 | 377,268 | 13,983 | 3,296 | 5,948 | 18,372 | 29,736 | 198 | 880 | 70,088 | 379,593 | 16 | 84 | 2c, 3, 3ka, 11, 17, 30, B, 10, 14a, 3bg, 14b | Kolmer et al 2013 |
| DURUM_09AZ103A | SRR630047 | DURUM | USA | 98 | 47 | 440,760 | 369,237 | 13,711 | 3,296 | 5,871 | 17,984 | 29,639 | 201 | 821 | 75,407 | 365,353 | 17 | 83 | 2c, 3, B, 10, 14b, 20 | Kolmer et al 2011 |
