## Supplemental table 1 for "Whole-genome sequencing of multiple isolates of *Puccinia triticina* reveals asexual lineages evolving by recurrent mutations"

Supplemental Table 2. Frequency of virulence to Thatcher lines of wheat near isogenic for single leaf rust resistance genes of *Puccinia triticina* isolates in groups from North America (NA) and Europe (EU), two groups virulent to durum wheat

| | Isolate Group | | | | | | | |
|---|---|---|---|---|---|---|---|---|
| | NA1 | NA2 | NA3 | NA4 | NA5 | NA6 | Durum | EU |
| **Lr1** | 0.500 | 0.818 | 1.000 | 1.000 | 0.906 | 1.000 | 0.000 | 0.077 |
| **Lr2a** | 0.000 | 0.000 | 0.143 | 1.000 | 0.875 | 0.333 | 0.000 | 0.077 |
| **Lr2c** | 0.250 | 1.000 | 0.143 | 1.000 | 0.875 | 1.000 | 0.077 | 0.923 |
| **Lr3** | 0.250 | 0.455 | 1.000 | 0.000 | 1.000 | 1.000 | 0.077 | 0.692 |
| **Lr9** | 0.000 | 0.182 | 0.114 | 0.000 | 0.281 | 0.000 | 0.000 | 0.000 |
| **Lr16** | 0.000 | 0.000 | 0.029 | 0.000 | 0.094 | 0.000 | 0.000 | 0.308 |
| **Lr24** | 0.000 | 0.091 | 0.286 | 0.000 | 0.594 | 0.000 | 0.000 | 0.000 |
| **Lr26** | 0.000 | 0.000 | 0.600 | 0.000 | 0.406 | 0.333 | 0.077 | 0.538 |
| **Lr3ka** | 0.000 | 0.455 | 0.514 | 0.000 | 0.188 | 0.333 | 0.000 | 0.692 |
| **Lr11** | 0.000 | 0.091 | 0.257 | 0.000 | 0.250 | 1.000 | 0.000 | 0.154 |
| **Lr17** | 0.250 | 0.000 | 1.000 | 1.000 | 0.000 | 1.000 | 0.000 | 0.615 |
| **Lr30** | 0.000 | 0.273 | 0.429 | 0.000 | 0.188 | 0.333 | 0.000 | 0.692 |
| **LrB** | 0.250 | 0.818 | 1.000 | 0.000 | 0.000 | 0.667 | 0.846 | 1.000 |
| **Lr10** | 0.500 | 0.909 | 0.886 | 0.500 | 0.969 | 0.667 | 1.000 | 0.923 |
| **Lr14a** | 1.000 | 0.455 | 1.000 | 0.000 | 0.656 | 1.000 | 0.000 | 0.538 |
| **Lr18** | 0.250 | 0.636 | 0.000 | 0.000 | 0.125 | 0.000 | 0.000 | 0.000 |
| **Lr3bg** | 0.250 | 0.000 | 1.000 | 0.000 | 0.031 | 0.333 | 0.000 | 0.692 |
| **Lr14b** | 0.750 | 1.000 | 0.971 | 1.000 | 1.000 | 0.000 | 0.923 | 0.923 |
| **Lr20** | 1.000 | 0.909 | 0.971 | 1.000 | 0.625 | 0.667 | 0.923 | 0.308 |
| **Lr28** | 0.500 | 1.000 | 0.000 | 1.000 | 1.000 | 0.667 | 0.000 | 0.000 |
| **Lr21** | 0.000 | 0.000 | 0.000 | 0.000 | 0.094 | 0.000 | 0.000 | 0.000 |
| **Lr39** | 0.000 | 0.182 | 0.171 | 0.000 | 0.344 | 0.000 | 0.923 | 0.000 |
| **Number of isolates** | 4 | 11 | 35 | 2 | 32 | 3 | 13 | 13 |
